# Shifts from pulled to pushed range expansions caused by reduction of landscape connectivity

**DOI:** 10.1101/2020.05.13.092775

**Authors:** Maxime Dahirel, Aline Bertin, Marjorie Haond, Aurélie Blin, Eric Lombaert, Vincent Calcagno, Simon Fellous, Ludovic Mailleret, Thibaut Malausa, Elodie Vercken

## Abstract

Range expansions are key processes shaping the distribution of species; their ecological and evolutionary dynamics have become especially relevant today, as human influence reshapes ecosystems worldwide. Many attempts to explain and predict range expansions assume, explicitly or implicitly, so-called “pulled” expansion dynamics, in which the low-density edge populations provide most of the “fuel” for the species advance. Some expansions, however, exhibit very different dynamics, with high-density populations behind the front “pushing” the expansion forward. These two types of expansions are predicted to have different effects on e.g. genetic diversity and habitat quality sensitivity. However, empirical studies are lacking due to the challenge of generating reliably pushed vs. pulled expansions in the laboratory, or discriminating them in the field. We here propose that manipulating the degree of connectivity among populations may prove a more generalizable way to create pushed expansions. We demonstrate this with individual-based simulations as well as replicated experimental range expansions (using the parasitoid wasp *Trichogramma brassicae* as model). By analyzing expansion velocities and neutral genetic diversity, we showed that reducing connectivity led to pushed dynamics. Low connectivity alone, i.e. without density-dependent dispersal, can only lead to “weakly pushed” expansions, where invasion speed conforms to pushed expectations, but the decline in genetic diversity does not. In empirical expansions however, low connectivity may in some cases also lead to adjustments to the dispersal-density function, recreating “classical” pushed expansions. In the current context of habitat loss and fragmentation, we need to better account for this relationship between connectivity and expansion regimes to successfully predict the ecological and evolutionary consequences of range expansions.

## Introduction

Range expansions and range shifts into novel habitats and landscapes are key ecological processes shaping the abundance and distribution of species (Sexton, McIntyre, Angert, & Rice, 2009). Understanding their ecological and evolutionary drivers and consequences has become especially relevant in the context of more frequent biological invasions (Miller et al., 2020; Renault, Laparie, McCauley, & Bonte, 2018) or increasing impacts of climate change (Hill, Griffiths, & Thomas, 2011; Lenoir et al., 2020).

Range expansions are usually modelled and analyzed in a framework based on the Fisher-KPP partial differential equation and its numerous declinations (see e.g. Hastings et al., 2005; Lewis, Petrovskii, & Potts, 2016). In this framework, range dynamics generally converge to solutions of constant profiles moving in space at a fixed velocity, called travelling waves. Travelling waves are considered “pulled” when spread is driven mostly, if not only, by the dynamics at the leading edge of the expansion. In biological range expansions, this happens when growth and dispersal rates are maximal at low densities; the velocity of the wave then only depends on low-density dispersal and growth (e.g. Birzu, Matin, Hallatschek, & Korolev, 2019). While growth and dispersal functions that are expected to generate pulled expansions certainly happen in nature (Harman, Goddard, Shivaji, & Cronin, 2020; Matthysen, 2005; Williams & Levine, 2018), they are not the only ones possible. For instance, growth rates can be reduced at low densities compared to intermediate ones, a phenomenon known as the Allee effect (Berec, Angulo, & Courchamp, 2007). Additionally, dispersal is often maximal at high densities (positive density-dependence) as it provides a mechanism to escape increased competition (Harman et al., 2020; Matthysen, 2005). In both cases, we would expect the advance of the expansion to be driven not by the low-density front populations, but by the population dynamics in a region located behind the front, where growth and/or dispersal are maximal. These “pushed” waves (Stokes, 1976) behave very differently than pulled waves (see e.g. Lewis et al., 2016). They will typically advance faster than expected based solely on growth and dispersal rates observed at low densities (i.e. in edge patches; Gandhi, Korolev, & Gore, 2019). The ratio of the actual expansion velocity to the velocity expected for the corresponding pulled expansion has been proposed as a quantitative indicator discriminating pulled, “semi-pushed” and pushed expansions (e.g. Birzu, Hallatschek, & Korolev, 2018). This ratio has been connected to other key metrics expected to vary along the pushed/pulled gradient (Birzu et al., 2018). In the context of biological range expansions or shifts, this includes the rate at which the edge or front of the expansion loses genetic diversity due to drift and successive founding events as it advances. Indeed, models and experiments show that neutral genetic diversity is lost at the expanding edge much more slowly in pushed than in pulled expansions (Birzu et al., 2018, 2019; Gandhi et al., 2019; Hallatschek & Nelson, 2008; Roques, Garnier, Hamel, & Klein, 2012), with estimated times to allele fixation differing by up to several orders of magnitude (Gandhi et al., 2019).

A key challenge for empirically studying the ecological and evolutionary consequences of pushed and pulled expansions (which, besides neutral genetic diversity, remain poorly known; see discussion in Birzu et al., 2019) is to obtain comparable expansions known to be either pushed or pulled. One could contrast strains, populations or closely related species naturally differing in density-dependent dispersal and/or Allee effects (Harman et al., 2020; Jacob, Chaine, Huet, Clobert, & Legrand, 2019; Matthysen, 2005; Walter, Grayson, Blackburn, Tobin, & Johnson, 2019). However, they are likely to also differ in other key traits (Jacob et al., 2019), making direct comparisons difficult. A better experimental solution would be to manipulate the focal species/population/strain’s environment to change the presence or strength of Allee effects and/or density-dependent dispersal. For instance, Gandhi et al. (2019; 2016) managed to reliably produce pushed expansions by changing the substrate on which *Saccharomyces cerevisiae* yeasts were grown: while yeasts grow best at low densities on galactose, they present an Allee effect when they have to use sucrose. Externally manipulating density-dependent dispersal may be more difficult, but could conceivably also be done by manipulating resource quality (Endriss et al., 2019; Van Allen & Bhavsar, 2014). Adding or removing conspecific cues independently of actual population size may also be an option (De Meester & Bonte, 2010), but not all organisms have resources or cues that are easy and useful to manipulate experimentally (Fellous, Duncan, Coulon, & Kaltz, 2012).

Rather than manipulating the within-habitat conditions, another solution that may be easier to generalize would be to reduce the level of connectivity among habitats. In a spatially structured environment, this could be done experimentally by e.g. altering the number, length or quality of the physical links between patches/populations. We hypothesize such experimental manipulations could lead to more pushed expansions because of dispersal stochasticity. Indeed, at low population sizes, reducing the dispersal rate sharply increases the risk the smaller populations at the edge of the expansion will fail to send any disperser, due to stochasticity alone (**Fig. 1**). Reduced dispersal rate could thus increase the influence of population density on dispersal success, leading to pushed expansions. This effect would occur when populations at the very edge of the expansions are of few individuals, typically less than a hundred (**Fig. 1**). Edge population densities low enough to trigger the effect described in **Fig. 1** are probably not uncommon in some taxonomic groups, where core population sizes are in the 100-1000 range (Krauss, Steffan-Dewenter, & Tscharntke, 2003; Santini, Isaac, & Ficetola, 2018). These are however much lower densities than those considered in most pushed expansion models and experiments (e.g. Birzu et al., 2018; Gandhi et al., 2019). The idea that expansions that should be pulled at asymptotically large carrying capacities can nonetheless appear pushed at lower densities, at least in terms of velocity dynamics, is not new. It was previously raised by Panja & van Saarloos (2002) and reviewed/expanded on in Panja (2004), which called these “weakly pushed” expansions (not to be confused with “semi-pushed” expansions *sensu* Birzu et al., 2018). “Weakly pushed” expansions arise when we stop assuming continuous fluid-like population densities can approximate the stochastic dynamics of discrete individuals in discrete/discretized space. Revisiting earlier range expansion studies including discreteness, we can indeed find hints that added dispersal stochasticity or decreased connectivity leads to population patterns resembling those seen in pushed expansions (Morel-Journel et al., 2016; Reluga, 2016; Williams & Levine, 2018). However, there is still no actual confirmation that reduced connectivity can produce pushed expansions (even in the “weakly pushed” sense), or that these expansions would share all key characteristics of pushed expansions that interest ecologists and evolutionary biologists (such as the dynamics of genetic diversity).

**Figure 1.**
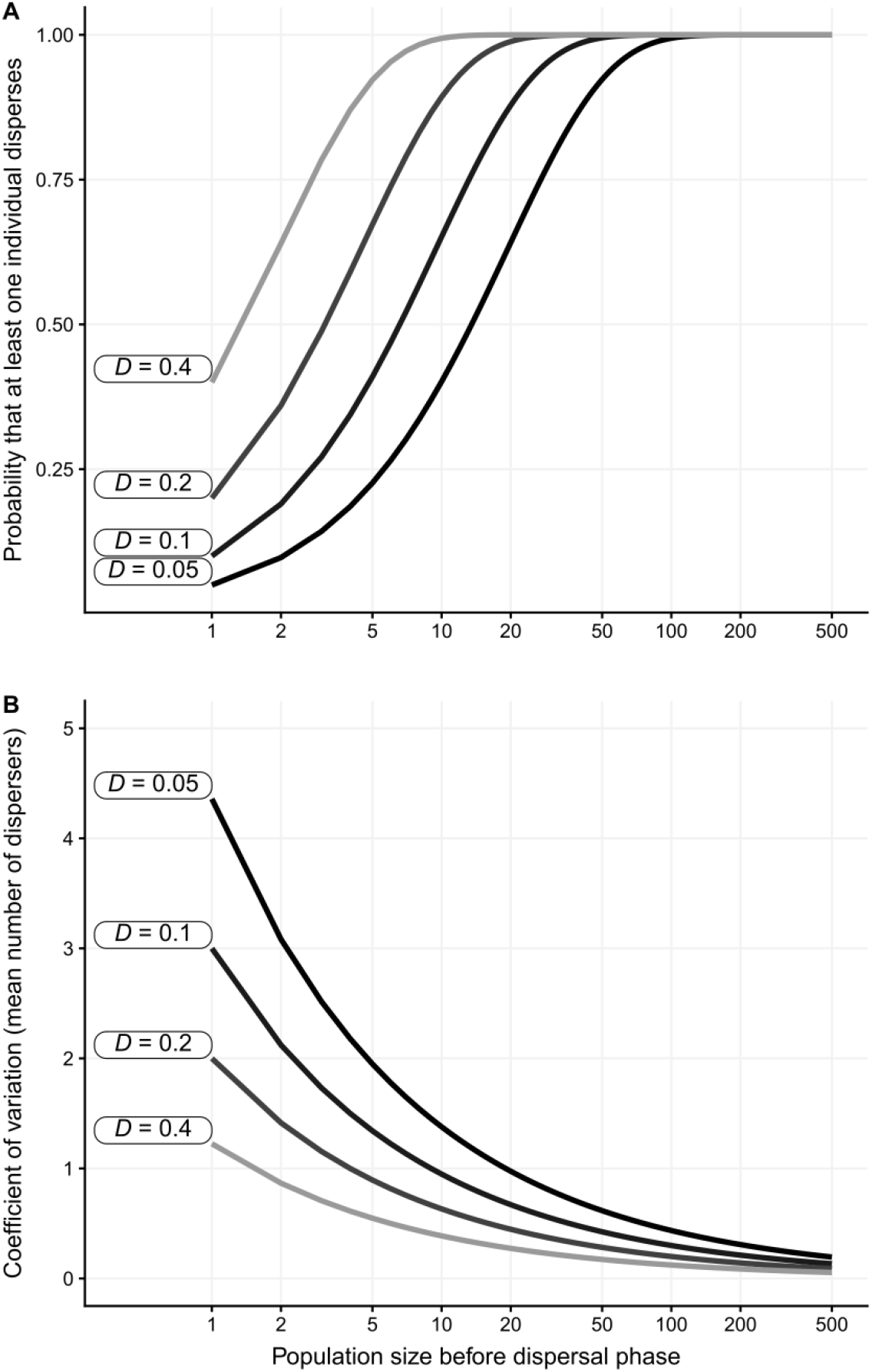
(A) Probability that at least one individual disperses from a patch (Pr(*n* > 0)) as a function of its population size *N* pre-dispersal and average dispersal rate *D*. (B) Coefficient of variation of the number of dispersers. Dispersal stochasticity is obtained by assuming the number of dispersers *n* from a patch is drawn from a Binomial distribution *n* ∼ Binom(*N*, *D*). Note the log scale on the x-axes.

In the present study, we combined simulations and experimental approaches to study the effect of reduced connectivity on the pushed vs. pulled status of expansions, at population sizes that are realistic for many “macroscopic” organisms. In individual-based simulations, we examined expansion velocities and the dynamics of neutral genetic diversity to (a) confirm Allee effects and density-dependent dispersal (Birzu et al., 2019) still lead to pushed expansions at “low” equilibrium population sizes *K*, and (b) show that reducing connectivity can also generate (weakly) pushed fronts even in the absence of the other two mechanisms. We complemented this approach by using minute wasps of the genus *Trichogramma* (Haond et al., 2018; Morel-Journel et al., 2016) in replicated experimental landscapes (Larsen & Hargreaves, 2020), to investigate whether reduced connectivity influenced the velocity and neutral genetic dynamics of “real” range expansions.

## Methods

### Simulations

To determine whether the new mechanism we propose, reduced connectivity, can actually generate pushed expansions, we used an individual-based model (IBM) approach. The model is in discrete time and space and simulates the dynamics of a haploid clonal species with non-overlapping generations, expanding in one direction in a one-dimensional landscape. Conceptually, the model is inspired by previous models by Birzu et al. (2018, 2019) and Haond et al. (2018), as well as Gandhi et al. (2016, 2019)’s experiments. Range expansions unfold during 100 generations in a landscape long enough that they never run out of empty patches. In practice, given the expansions advance in only one direction and individuals can only disperse at most one patch per generation (see below), this means any landscape length higher than the number of generations. All patches are of equal and constant quality, determined by the equilibrium population density *K*.

At the start of the expansion, *K* adult individuals that have not yet reproduced are introduced in one patch at one of the two extremities of the landscape (coordinate *x* = 0). To be able to later study genetic diversity, all individuals are randomly assigned one of two allele values (coded 0 and 1) at a neutral genetic locus *L*. Each generation until the end of a run, the life cycle then happens as follows:

1. Adult individuals disperse with a probability *D* = logit^−1^(logit(*D*_0_) + *β*_*density*_ × *N*/*K*), where *N* is the patch population size immediately before the dispersal phase, *D*_0_ is the (hypothetical) dispersal rate at *N* = 0, and *β*_*density*_ the slope of the dispersal-density relationship on the logit scale. This function is based on the way empirical dispersal-density data are usually analyzed (through generalized linear models; see e.g. De Meester & Bonte, 2010; Van Allen & Bhavsar, 2014). Dispersers randomly move to one of the nearest neighboring patches; that is, the maximal dispersal distance is of 1 patch.
2. Reproduction occurs post-dispersal; the number of offspring *F* each individual produces is drawn from a Poisson(*μ*) distribution. The mean fecundity *μ* is based on a Ricker equation modified to allow potential Allee effects (Morel-Journel et al., 2016): 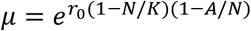. In this equation, *r*_0_ is the hypothetical population growth rate at *N* = 0 in the absence of Allee effects, and *A* an Allee threshold such that *A* = 0 leads to no Allee effects, 0 < *A* ≤ 1 leads to weak Allee effects (*sensu* e.g. Berec et al., 2007, i.e. where positive density-dependent growth never leads to negative growth rates), and *A* > 1 leads to strong Allee effects (where growth rates are negative for *N* < *A*). All new individuals inherit the value at the neutral locus *L* from their parent with no mutation.
3. All adults die; juveniles then become adults.

The model was written in Netlogo (Wilensky, 1999), version 6.1.1, and set up using the *nlrx* R package (Salecker, Sciaini, Meyer, & Wiegand, 2019). We tested 5 scenarios (× 2 possible values of *K*; see below and **Fig. 2**). The “reference” scenario had no Allee effects, no density-dependent dispersal (*β*_*density*_ = 0) and a dispersal rate *D*_0_ set to 0.2, a biologically “typical” rate according to experimental (e.g. Fronhofer et al., 2018) or natural observations (Marjamäki, Contasti, Coulson, & McLoughlin, 2013; Stevens et al., 2013), spanning many taxa. Three other scenarios, which we expected to lead to pushed expansions, each differed from the reference by one parameter: either a weak Allee effect was present (*A* = 0.95), there was positive-density-dependent dispersal (*β*_*density*_ = 1) or connectivity was reduced by half (*D*_0_ = 0.1). The fifth scenario was a combination of reduced connectivity and density-dependent dispersal (*β*_*density*_ = 1 and *D*_0_ = 0.1). In all cases *r*_0_ was set to log(5), which is well within the range of plausible values for insects (Hassell, Lawton, & May, 1976). We tested each scenario for two values of the equilibrium density *K*, 225 and 450 individuals per patch, as the relationship between *K* and the expansion velocity *v* is expected to differ between pushed and pulled expansions (Haond et al., 2018). These densities are within the range of *K* used in Haond et al. (2018)‘s simulations and experiments, and correspond respectively to 50% and 100% of the largest possible population in our own experimental expansions (see below). These values of *K* are also within one order of magnitude of those seen for some species of butterflies (Krauss et al., 2003) or herbaceous plants (Dauber et al., 2010) in patchy environments, or in land mollusks (Kappes et al., 2009) assuming a typical home range of a few m² at most (Bailey, 1989). Each scenario × *K* combination was replicated 100 times.

**Figure 2.**
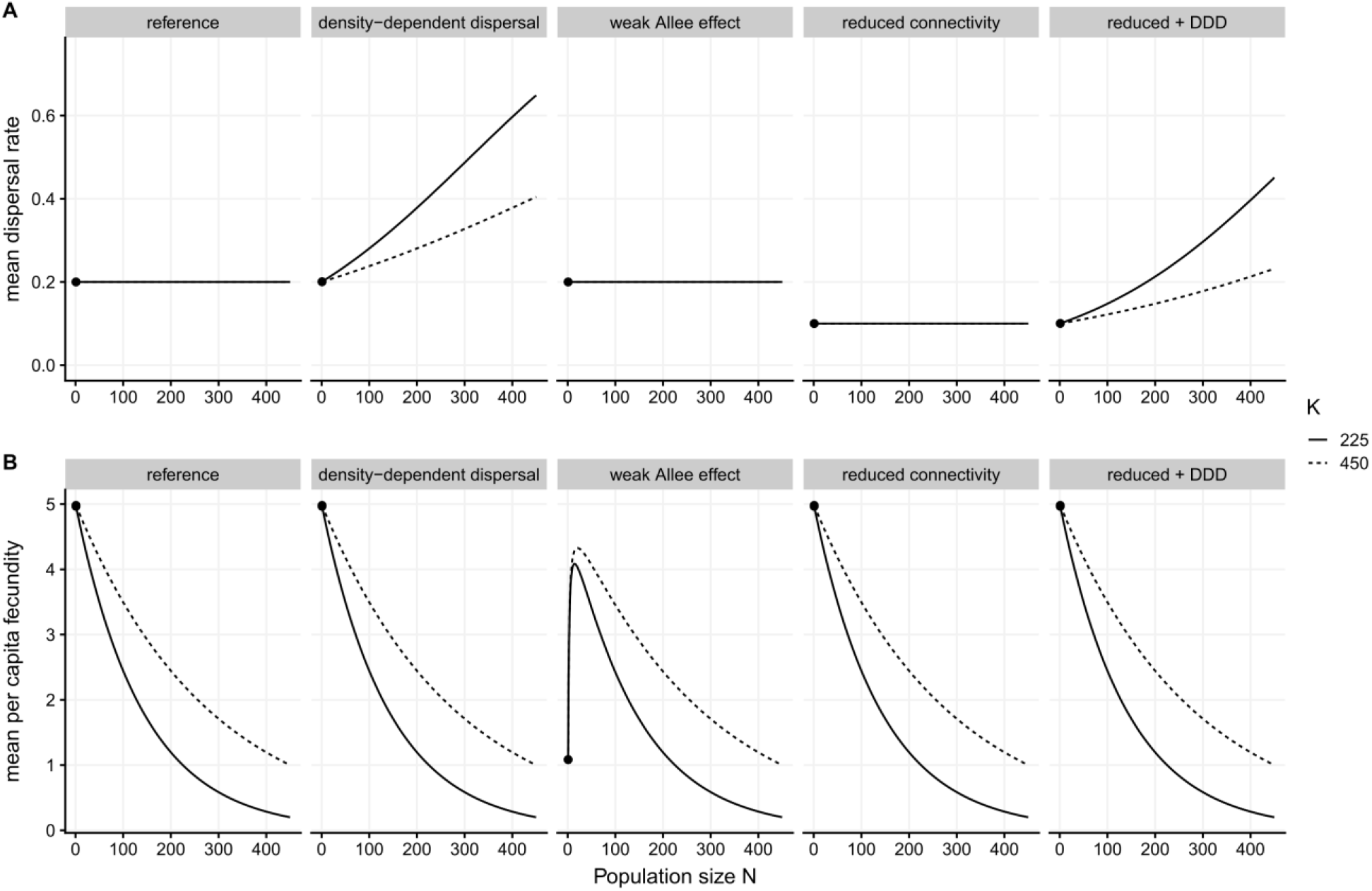
Dispersal (A) and fecundity (B) functions used in simulations. The functions differ depending on equilibrium population size *K* because they are defined in terms of *N*/*K*, rather than *N*. Dots mark the values of dispersal and fecundity used in the definition of the reference velocity *v*_*F*_.

### Experimental range expansions

We used laboratory strains of the haplo-diploid egg parasitoid *Trichogramma brassicae* Bezdenko, 1968 (Hymenoptera: Trichogrammatidae) as our model in experiments (**Supplementary Material 1**). *T. brassicae* wasps are raised routinely in the lab using eggs of the Mediterranean flour moth *Ephestia kuehniella* Zeller 1879 (Lepidoptera: Pyralidae) as substitution host. *E. kuehniella* eggs are irradiated before use, which prevents their larval development while still allowing *Trichogramma* wasps to use them as hosts (St-Onge, Cormier, Todorova, & Lucas, 2014). To be able to better generalize our results, we used three independent genetic mixes of *T. brassicae* (**Supplementary Material 1**). Note that the effect of genetic background itself is not the object of the present manuscript, and is thus not analyzed directly in any of our statistical models, for simplicity (as in Van Petegem et al., 2018).

To experimentally test the effects of reduced connectivity on range expansions, we set up a series of one-dimensional artificial landscapes (**Fig. 3**), in which we monitored *T. brassicae* wasps for 14 non-overlapping generations (including initially released adults, i.e. generation 0). The landscapes were made of closed plastic vials (5 cm diameter, 10 cm height) connected to each of their nearest neighbors by either three 20 cm flexible tubes (“reference”-level connectivity) or one 40 cm tube (reduced connectivity) (tube internal diameter: 5 mm). Landscapes were initially 5 patches long (including the release patch), to allow dispersal kernels with movements > 1 patch, and extended as needed to ensure there were always at least 2 available empty patches beyond the current front. Each treatment was replicated 12 times (4 per genetic mix), for a total of 24 experimental landscapes. Landscapes were initiated by introducing ≈ 300 adult wasps (range: 261–324) in one extremity patch (*x* = 0), so the expansion could only advance in one direction in each landscape. Landscapes were kept under standardized conditions (23°C, 70% relative humidity, 16:8 L:D). Each generation then unfolded along the following steps:

- (a) We provided approximately 450 new eggs of *Ephestia kuehniella* per patch for wasps to lay eggs in. *E. kuehniella* eggs were presented on paper strips to facilitate handling.
- (b) Adults (and old egg strips) were removed after 48 hours to enforce non-overlapping generations and standardized generation times.
- (c) When parasitoid larval development was advanced enough to identify signs of parasitism (host eggs darkening after ≈ 7 days), we temporarily removed egg strips from patches and assessed the presence/ absence of parasitized eggs by eye. We photographed all patches with parasitized eggs for semi-automatic estimations of population sizes (Nikon D750 60mm f/2.8D macro lens, 24.3 Mpix). Population sizes were estimated (as % of host eggs parasitized) using ImageJ and the Codicount plugin (Abramoff, Magalhães, & Ram, 2004; Perez, Burte, Baron, & Calcagno, 2017). We used four different macros each trained on a different set of pictures, to account for “observer” effects; each macro tended to give consistently biased estimates, but combining them led to on average unbiased estimates (**Supplementary Material 2**). These population size estimates should be relatively robust to superparasitism, as in the vast majority of cases only 1 wasp per host survives to this stage, even when multiple eggs have been laid in the host (Corrigan, Laing, & Zubricky, 1995). Eggs were then replaced in their patches until the emergence of the adults, when we started a new cycle from step (a).

**Figure 3.**
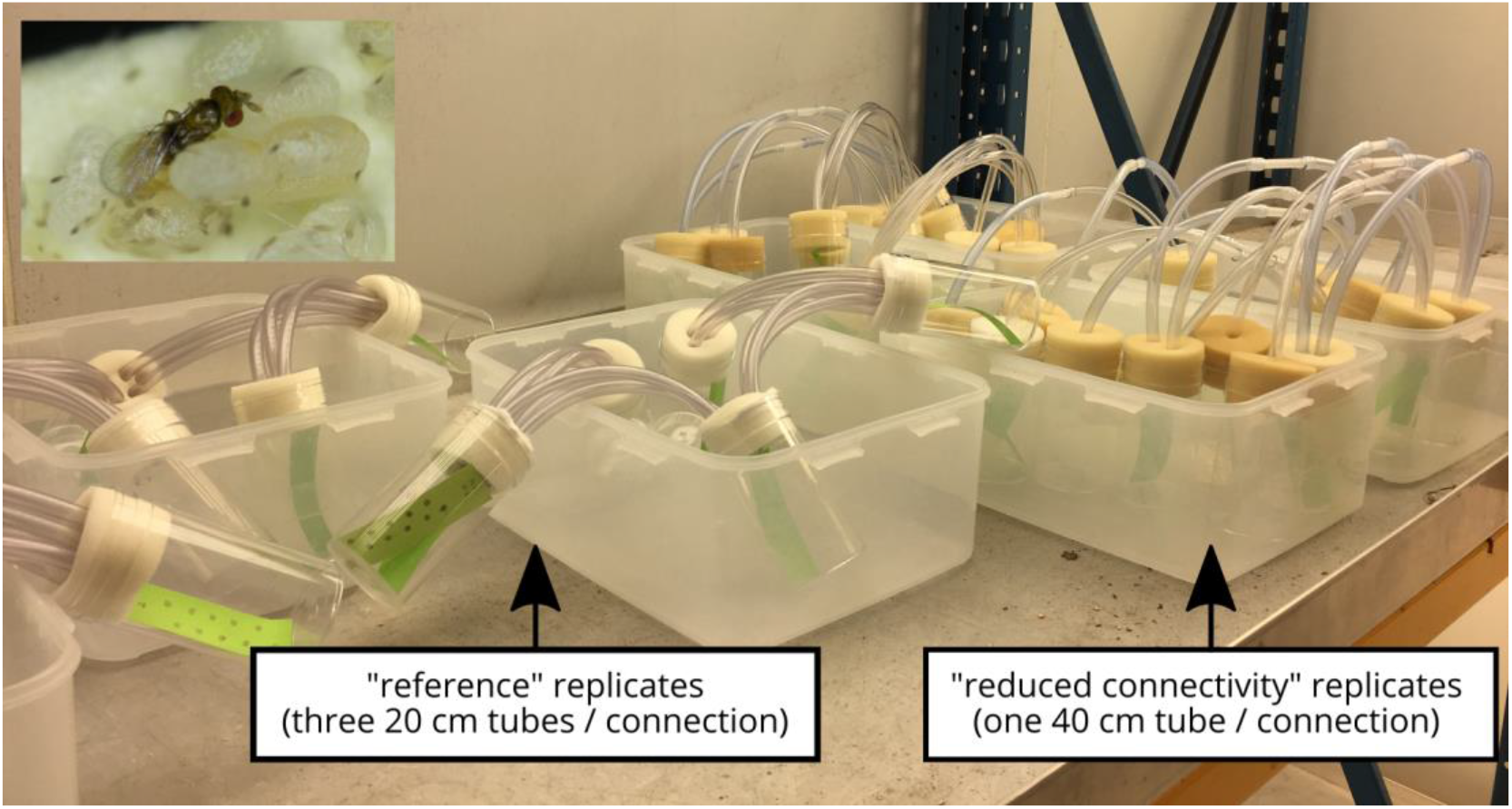
A representative subset of the experimental landscapes, showing the “patches” connected by 1 or 3 tubes, depending on treatment, as well as clusters of host eggs on paper strips for easy manipulation. Picture by Aline Bertin. Inset: *Trichogramma brassicae* on *Ephestia kuehniella* eggs. Picture by Géraldine Groussier, used with authorization.

We confirmed the experimental design led to reduced connectivity by comparing the average egg-laying distances (in patches from release sites) between treatments at the start of the experiment, i.e. the only time point when the source patch of all egg-laying individuals was known. As dispersal is defined as “movement of individuals or propagules with potential consequences for gene flow across space” (Ronce, 2007), this metric is in effect a measure of average (effective) dispersal distance. The average distance was reduced by ≈ 20% in “reduced connectivity” landscapes compared to “reference” ones (**Supplementary Material 3**).

To determine how genetic diversity evolved during experimental range expansions and whether it was influenced by connectivity, we kept and genotyped adult female wasps after their removal from the experimental landscapes. In each landscape, we sampled initially released individuals (i.e. generation 0; hereafter “origin” wasps) as well as wasps found in both the release patch (“core” wasps) and the most advanced population (“edge” wasps) at generations 4, 8 and 12. In each patch × time combination, we aimed to sample about 20 individuals; the actual number was sometimes lower especially in edge populations with fewer individuals (average ± SD: 18.10 ± 2.63 wasps; range: 4-28; total: 3043). Wasps were genotyped at 19 microsatellite loci; microsatellite characteristics, as well as the DNA extraction and amplification protocols, are detailed in **Supplementary Material 4.**

This experiment complied with all relevant national and international laws; no ethical board recommendation or administrative authorization was needed to work on or sample *Trichogramma brassicae*.

### Statistical analyses

All data (experimental and simulated) were analyzed in a Bayesian framework, using R, version 4.0.3 (R Core Team, 2020) and the *brms* package version 2.14.4 (Bürkner, 2017) as frontends to the Stan language (RStan version 2.19.3; Carpenter et al., 2017; Stan Development Team, 2018). We used (non)linear multilevel/mixed models. The model descriptions below are summaries aiming for clarity rather than completeness; formal and complete write-ups for all statistical models are given in **Supplementary Material 5**. We use general-purpose “weakly informative” priors based on McElreath (2020) for all parameters except one where there is strong theoretical prior knowledge (see **Supplementary Material 5**). For each model, we ran four chains with enough iterations to ensure effective sizes were satisfactory for all fixed effect, random effect and distributional parameters (both bulk- and tail-ESS *sensu* Vehtari, Gelman, Simpson, Carpenter, & Bürkner, 2020 > 1000). In addition to graphical posterior checks (Gabry, Simpson, Vehtari, Betancourt, & Gelman, 2019; Vehtari et al., 2020), we checked chain convergence using the improved 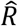 statistic by Vehtari et al. (2020). All credible intervals given in text and figures are 95% Higher posterior Density Intervals (HDIs). Data handling and figure generation were mainly done using the various *tidyverse* packages (Wickham et al., 2019), as well as the *tidybayes* (Kay, 2019) and *patchwork* (Pedersen, 2019) packages.

#### Expansion velocity (simulations and experimental data)

In the long run, the position *X*_*t*_ of the front (here using the distance between the farthest populated patch and the release site) is expected to increase linearly with time *t*, whether expansions are pushed or pulled (e.g. Lewis et al., 2016):

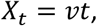

where *v* is the asymptotic expansion velocity. However, expansions only settle on the velocity *v* after a transition period; the above equation is likely ill-suited for estimating front positions when expansions are followed during their early stages and population sizes are low (so stochasticity is high). In these cases, which correspond to the present study, we propose to use the following model:

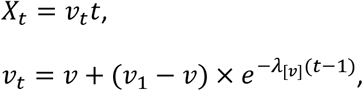

where *v*_1_ is the initial velocity after 1 generation of expansion (we use *v*_1_ as our “starting point”, and thus *t* – 1, because estimating velocities at *t* = 0 leads to convergence problems), and *λ*_[*v*]_ the rate of exponential convergence to the asymptotic velocity *v*.

We fitted this model to both simulated and experimental front locations, assuming a lognormal distribution to account for the fact distances travelled are always strictly positive. We note that one could assume a power-law decay, rather than an exponential one (Panja, 2004). We found however that a power-law model performed either much worse (simulated data) or similarly (experimental data) than the exponential one, and thus used the exponential decay in both cases (see **Supplementary Material 6**, as well as the companion script for detailed model comparisons, **Data availability**).

For experimental data, the submodels for log(*v*_1_), log(*v*) and log10(*λ*_[*v*]_) included fixed effects for treatment as well as random effects of replicate identity. For simulated data, we made three slight adjustments. First, we only used data from every fifth generation, to reduce computation time (by about an order of magnitude, based on preliminary tests) without significant impact on predictive success. Second, the submodel for *v* used logit(*v*)instead of log(*v*), as velocities were by design ≤ 1 (due to nearest neighbour dispersal). Finally, the model was simplified by setting *v*_1_ to 1; given the initial population size at *t* = 0, all simulated landscapes were all but guaranteed to send at least one individual to the next patch during the first generation, (see **Fig. 1**), leading to a velocity ≈ 1 patch/generation at *t* = 1.

In simulations, the growth and dispersal functions are fully known (**Fig. 2**), so we were also able to directly compare the estimated *v* to their respective *v*_*F*_, the predicted velocity for a pulled wave with the same *D*_0_ and *r*_0_ (Birzu et al., 2019; Lewis et al., 2016):

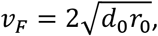

where *d*_0_ = 0.5*D*_0_ for a one-dimensional landscape with nearest neighbour dispersal (see e.g. Haond et al., 2018). Pulled expansions are expected to have 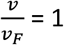, fully pushed expansions to have 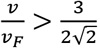, and so-called semi-pushed expansions to be in-between (Birzu et al., 2018, 2019). Because the formula we use for Allee effects leads to a fecundity of 0 at density = 0, and because growth and dispersal at density = 0 are not meaningful anyway, we used *r*_1_ and *D*_1_ at *N* = 1 to estimate *v*_*F*_ rather than the “true” *r*_0_ and *D*_0_. Note that the formula we use for *v*_*F*_ is based on the continuum assumption, and should not be exactly valid for discrete systems with stochasticity in growth, like ours. In fact, we would expect *v*_*F*_ < *v*_*F*[continuous]_ for such systems (e.g. Hallatschek & Korolev, 2009). However, some of us previously showed that for a simulated system like ours, using the simpler continuum formula for *v*_*F*_ is good enough, with an expected bias of ≈ 1% for 200 ≲ *K* ≲ 500 (Haond et al., 2018). This is much less than the smallest difference in speeds between pulled and fully pushed expansions (≈ 6%, because 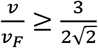), and should therefore not affect our decision to classify a type of expansion as pushed or pulled.

#### Relationship between equilibrium population density K and velocity v

It has been demonstrated that, over the range of densities we consider in our experiments and simulations, the correlation between *K* and the asymptotic velocity *v* could be another indicator of pushed expansions (Haond et al., 2018). Pushed expansions would show a positive correlation between *K* and *v*, while pulled expansions should in theory show no correlation. However, this indicator cannot be considered alone, as demographic stochasticity can also create such a correlation (Brunet & Derrida, 1997; Hallatschek & Korolev, 2009). The main difference seems to be that *K*-*v* correlations created by demographic stochasticity alone are associated with 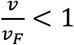 (Brunet & Derrida, 1997; Hallatschek & Korolev, 2009), while 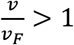 pushed expansions (see above).

In simulations, we simply compared the posterior distributions of *v* at *K* = 225 and *K* = 450, as *K* was fixed.

In experiments, we used the fact that *K* and *v* spontaneously varied among replicates, and analyzed among-replicate correlations using a bivariate multilevel model. This allowed us to estimate correlations accurately while accounting for various sources of uncertainty, including observer/macro error for population sizes (**Supplementary Material 5**). We first started by designing univariate models. The univariate model for velocities *v* is the one described in the previous section. For *K*, we used estimated population sizes in the starting patch (*x* = 0), as this was the only patch we expected to be at equilibrium from the start, based on release densities (Morel-Journel et al., 2016). We initially assumed the estimated percentage of hosts identified as “parasitized” could be analyzed using a Beta model, with fixed effects of experimental treatment and random effects of replicate landscape and computer macro, the latter to account for consistent macro-level bias (**Supplementary Material 5**). However, we found using posterior checks (Gabry et al., 2019) that this model failed to accurately represent data distribution and variability. We then used instead a Student *t* model on logit-transformed percentages, with the same fixed and random effects; this performed better (see companion code, **Data availability**).

Following this, we fitted a bivariate model using the two selected models for front position and population sizes. Replicate-level random effects for front parameters and for *K* were combined in the same variance-covariance matrix; this covariance matrix was estimated separately for each treatment. This allowed us to obtain treatment-specific posteriors for the replicate-level correlation between *K* and *v*.

#### Genetic diversity, experimental data

When expected heterozygosity (Nei, 1973) *H* is used as a measure of genetic diversity, theory predicts the genetic diversity both in core patches and at the edge of a range expansion should decay exponentially with time *t* (Birzu et al., 2018; Hallatschek & Nelson, 2008):

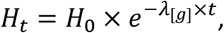

where *H*_0_ is the initial heterozygosity, and *λ*_[*g*]_ the rate of decay of genetic diversity. The decay rate *λ*_[*g*]_ is proportional to the inverse of the effective population size *N*_*e*_, with the exact relationship depending on ploidy (see e.g. Coop, 2020). For experimental data, we used this equation directly in a non-linear model to estimate whether the dynamics of genetic diversity varied between our two treatments and between core and edge patches. Multilocus expected heterozygosity was first calculated for each location (core/edge) × time combination using microsatellite data and the *adegenet* package (Jombart, 2008). Our submodel for *λ*_[*g*]_ included fixed effects for treatment, location (initial release patch for “core”/ most advanced patch for “edge”) and their interactions, as well as a random effect of replicate identity. The submodel for the initial diversity *H*_0_ only included the random effect of replicate, as we did not expect differences between treatments beyond random fluctuations, and core/edge patches are the same at *t* = 0. Location was included in the *λ*_[*g*]_ submodel as a centred dummy variable (−0.5 for “core”, 0.5 for “edge”) set to 0 at *t* = 0 as, again, core/edge patches are the same at *t* = 0. The submodels were estimated on logit(*H*_0_) and log10(*λ*_[*g*]_) to keep them within proper bounds; expected heterozygosities are proportions, and the decay rate *λ*_[*g*]_ must be positive. We fitted the overall model on logit(*H*_*t*_) using a Student *t* distribution, rather than on *H*_*t*_ using a Beta distribution (and a logit link). This is because the former is likely to be more robust to rare outliers (Kruschke, 2015), like those caused by sampling effects before genotyping (see companion script for detailed model comparisons, **Data availability**).

#### Genetic diversity, simulated data

With simulated data, we cannot use the equation above as our basis to fit a model. This is because the way we simulate neutral genetic diversity (one locus with only two alleles in a haploid species) means values of *H*_*t*_ = 0 are very frequent (especially as *t* increases), and our previous model cannot handle data containing zeroes. We instead used the fact that, with two alleles and if a given treatment is replicated a large number of times, a version of *λ*_[*g*]_ can also be recovered from the way among-replicate variance in allelic frequencies *V* (for either allele) changes with time (e.g. Gandhi et al., 2019):

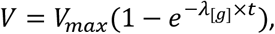

where *V*_*max*_ is the asymptotic variance reached when all replicates have fixed one of the alleles, which is equal to the product of initial allelic frequencies. As for experimental data, our submodel for log10(*λ*_[*g*]_) included fixed effects for treatment, location (core/edge) and their interactions. The submodel for logit(*V*_*max*_) included only a constant intercept, as *V*_*max*_ should be identical between all cases (and ≈ 0.25) but for random sampling fluctuations. We fitted this model on all data with *t* > 0 using a Beta distribution, as the issues raised for experimental data above regarding outliers did not apply (each data point here being the summarized outcome of 100 independent populations).

## Results

### Expansion velocity *v* and correlation with *K*

In simulations, absolute asymptotic expansion velocity *v* differed between treatments. Density-dependent dispersal, when alone, led to higher velocities than in reference expansions, while Allee effects and reduced connectivity (with or without density-dependent dispersal) led to slower expansions (**Fig. 4**, **Supplementary Materials 6,7**). Combining reduced connectivity and density-dependent dispersal led to velocities closer to the reference than reduced connectivity alone (**Fig. 4**, **Supplementary Materials 6,7**). Relative velocities 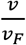 in the reference landscapes were very close to those expected for pulled expansions (1.01 and 0.99 for *K* = 450 and 225 respectively, **Fig. 4**). The other four treatments all had higher velocity ratios and were firmly in the range corresponding to true pushed expansions based on Birzu et al. (2018) (average velocity ratios 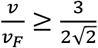, **Fig. 4**). Expansions were faster when *K* = 450 than when *K* = 225 for all five treatments (**Fig. 4, Supplementary Materials 6,7**).

**Figure 4.**
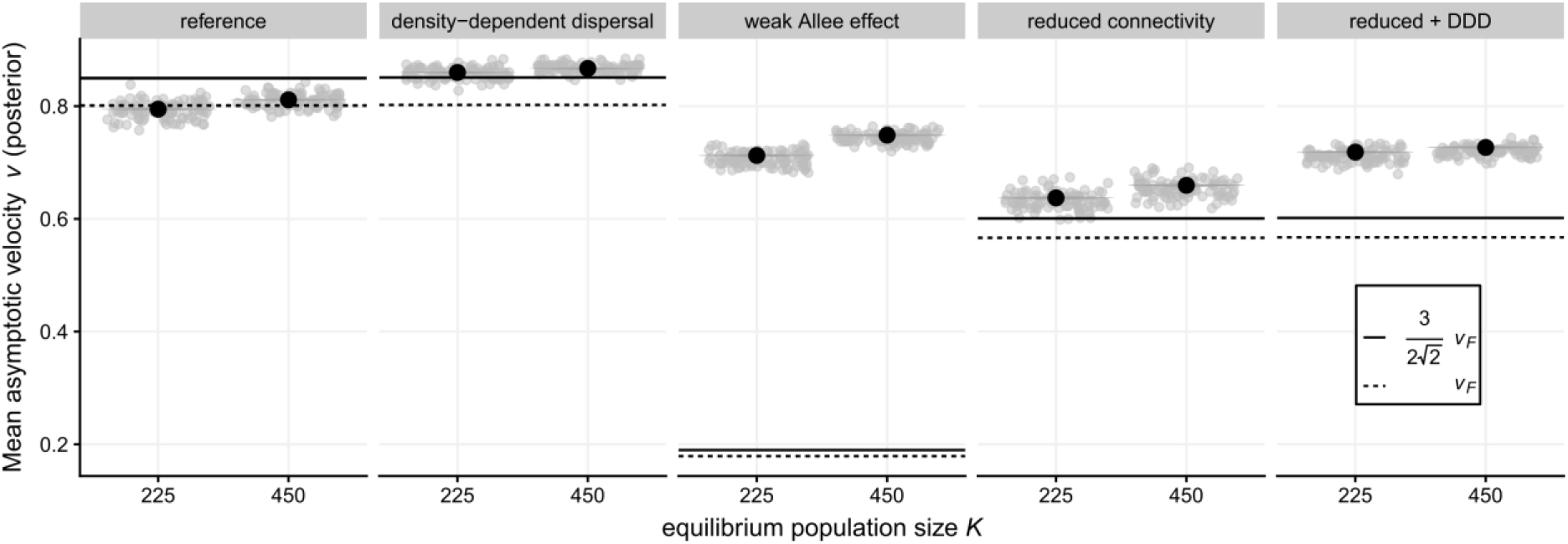
Posterior distribution of the average asymptotic expansion velocity (*v*) in simulations, depending on simulation scenario and equilibrium population size *K* (black dots: posterior means, the posterior is in all cases narrower than the dot used to depict the mean). The horizontal lines mark the range between 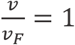 (i.e. pulled expansions) and 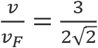 (limit between semi-pushed and fully pushed expansions; Birzu et al., 2018, 2019). Light grey dots are observed speeds at the end of each simulation run (*t* = 100).

In experiments, absolute estimated asymptotic velocities were virtually indistinguishable between reference and “reduced connectivity” landscapes (**Fig. 5**). There was no correlation between replicate-level estimates of *K* and *v* in “reference” landscapes (*r* = 0.04 [−0.47; 0.57], **Fig. 6**), while there was a positive correlation when connectivity was reduced (*r* = 0.51 [0.02; 0.92], **Fig. 6**). One must note however that credible intervals are wide, meaning we cannot say with certainty that the two correlations are different (Δ = 0.47 [−0.22; 1.20]). There was also no clear evidence that mean population sizes in the “core” starting patch differed between treatments, with 59.6% [44.6; 72.7] of hosts successfully parasitized on average in “reference” landscapes vs 63.7% [50.4; 77.0] in “reduced connectivity” landscapes (based on univariate model; Δ = −4.07% [−9.36; 1.39])(**Fig. 6A**).

**Figure 5.**
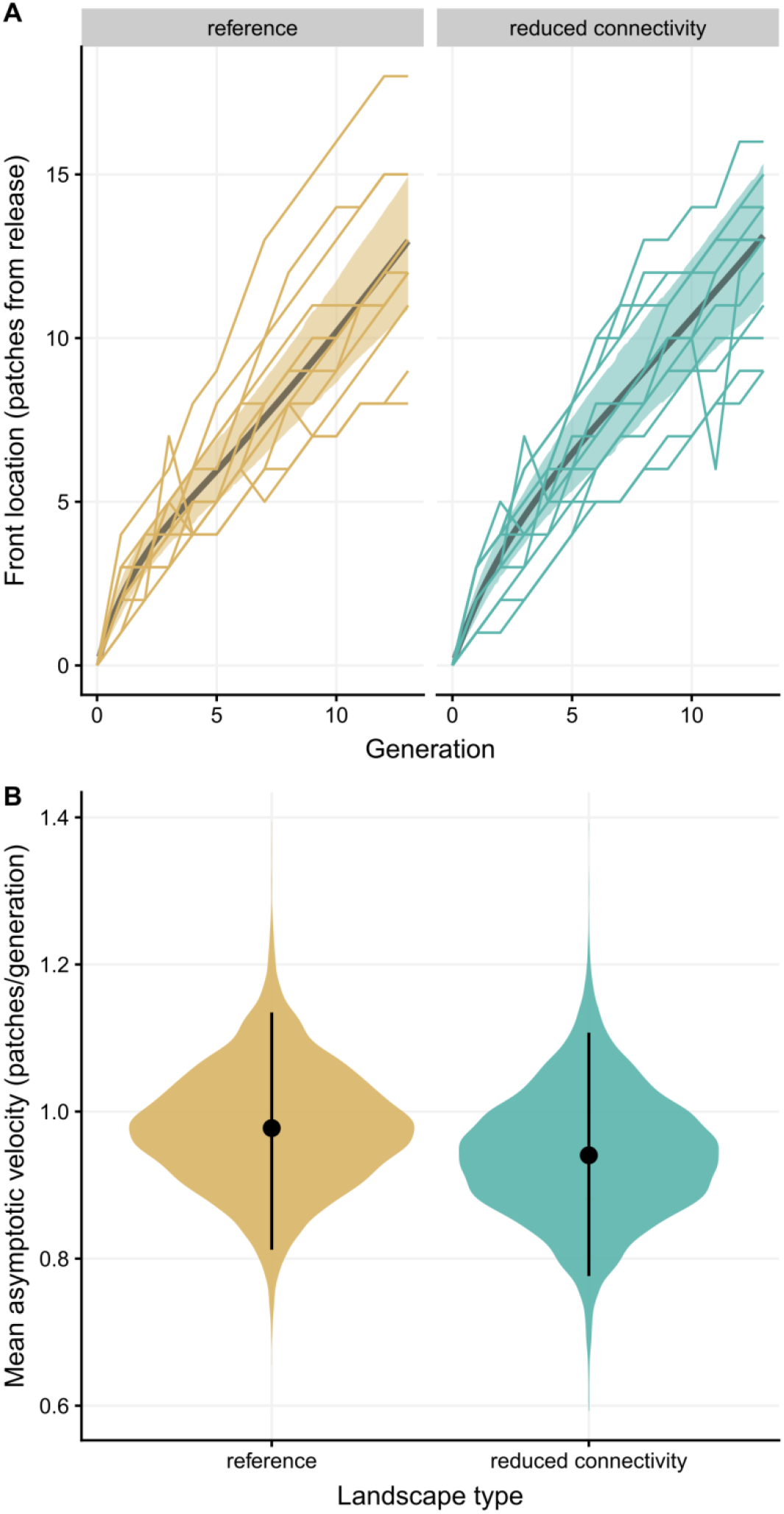
(A) Front locations as a function of the number of generations since release and experimental landscape type. Posterior means and 95% credible bands are displayed along observed trajectories for each landscape. (B) Posterior average asymptotic velocity *v* as a function of landscape type. Dots are posterior means, vertical bars 95% credible intervals.

**Figure 6.**
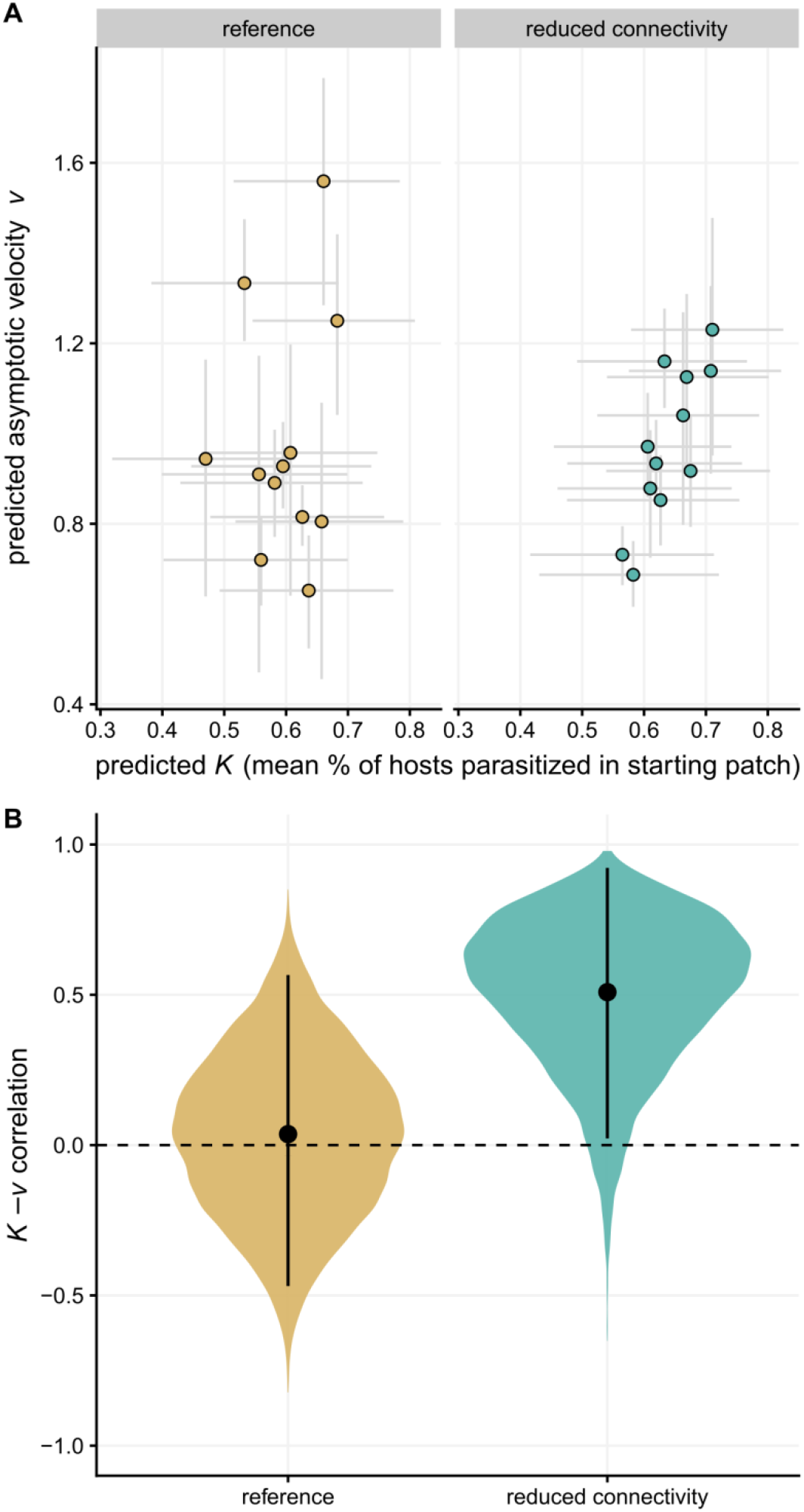
Correlation between population size in the core *K* and asymptotic velocity *v* in experiments. (A) Replicate-level predictions for *K* and *v* as a function of treatment (see text for comparison of means). (B) Posterior distribution of the *K*-*v* correlation coefficient (latent-scale) as a function of treatment. In both subplots dots are posterior means, bars 95% credible intervals.

### Shifts in genetic diversity

In simulations, genetic diversity declined faster, as measured by the decay rate *λ*, in edge patches than in core patches for all treatments (**Fig. 7**, **Supplementary Materials 6,7**). In edge patches, the addition of density-dependent dispersal and Allee effects led to a slower rate of diversity loss compared to the reference (**Fig. 7**, **Supplementary Materials 6,7**). By contrast, reducing connectivity led to a faster rate of diversity loss (**Fig. 7**, **Supplementary Materials 6,7**), but only when there was no density-dependent dispersal at the same time. Core patch dynamics were much less variable. However, when a treatment did deviate from the reference, it mostly did so in the same direction as in edge patches (see e.g. “density-dependent dispersal” or “reduced connectivity” at *K* = 225, or “reduced connectivity + density dependent dispersal” at *K* = 450; **Fig. 7**, **Supplementary Materials 6,7**). In all five treatments and whether we looked at core or edge patches, genetic diversity was lost faster when *K* = 225 than when *K* = 450 (**Fig. 7**, **Supplementary Materials 6,7**).

**Figure 7.**
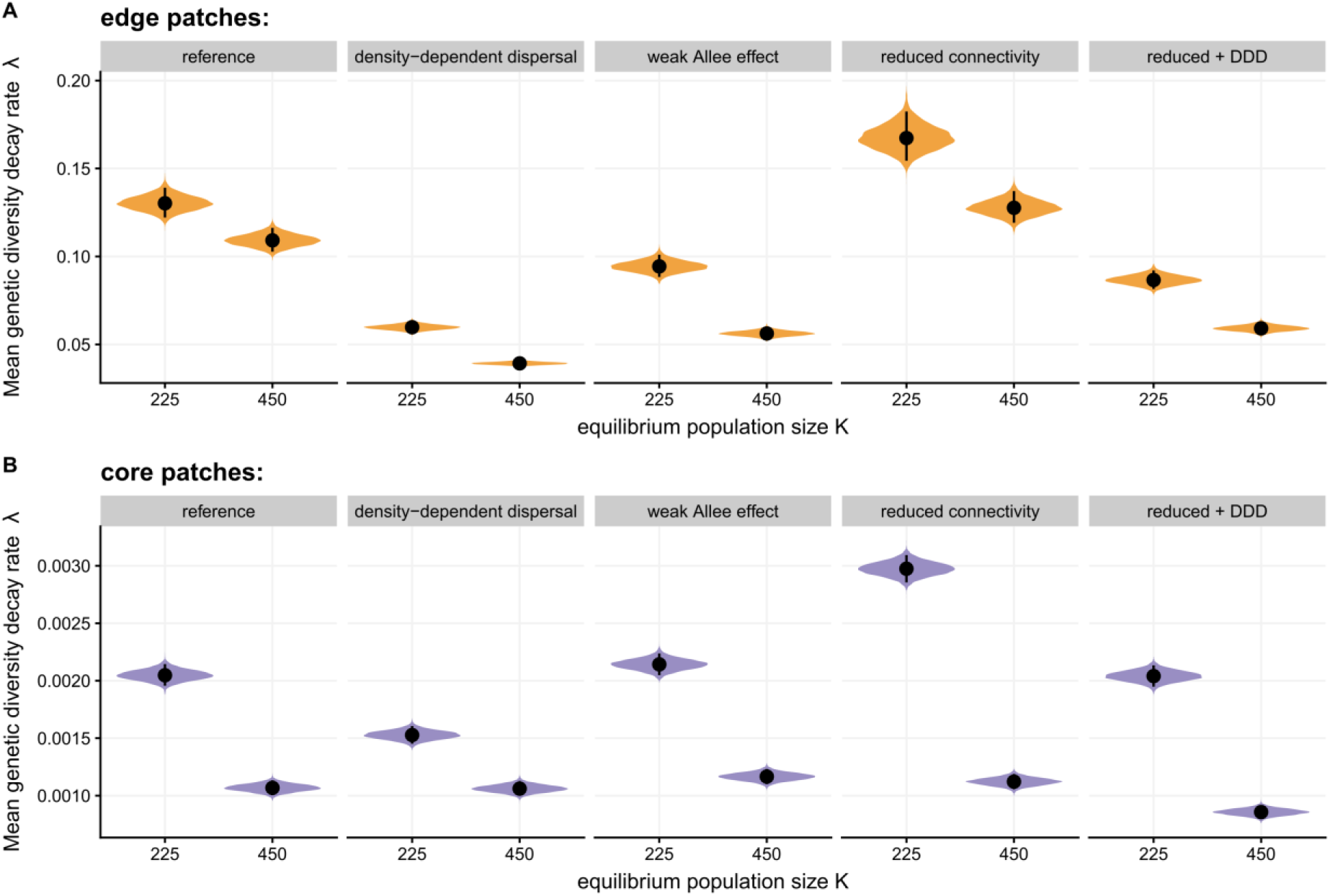
Posterior distribution of the average decay rate of genetic diversity with time (*λ*) in simulations, depending on simulation scenario, equilibrium population size *K* and patch location (either the most advanced patch at the time of measure (edge, A), or the original release site (core, B)). Dots are posterior means, vertical bars 95% credible intervals. Please note that posteriors for core and edge patches are displayed on different scales on the y-axis, for readability.

In the experimental landscapes, genetic diversity also decayed on average in all tested contexts of our experiment (**Fig. 8**), and we also found differences between edge and core patches and between both treatments. The rate of genetic diversity loss did not differ clearly between treatments in core patches (mean Δ_reference−reduced_ = - 6.5 × 10^−3^, 95% CI : [−15.5 × 10^−3^, 2.1 × 10^−3^]), but diversity was lost more rapidly in reference edge patches than in “reduced connectivity” ones (mean Δ_reference−reduced_ = 18.3 × 10^−3^, 95% CI : [1.8 × 10^−3^, 35.1 × 10^−3^]) (**Fig. 8**, **Supplementary Material 7**). In addition, while edge patches lost diversity faster than core patches in reference landscapes, (mean Δ_edge−core_ = 40.6 × 10^−3^, 95% CI : [27.7 × 10^−3^, 53.7 × 10^−3^]), this difference was still present but reduced when connectivity was reduced (mean Δ_edge−core_ = 15.8 × 10^−3^, 95% CI : [8.0 × 10^−3^, 24.3 × 10^−3^]) (**Fig. 8)**.

**Figure 8.**
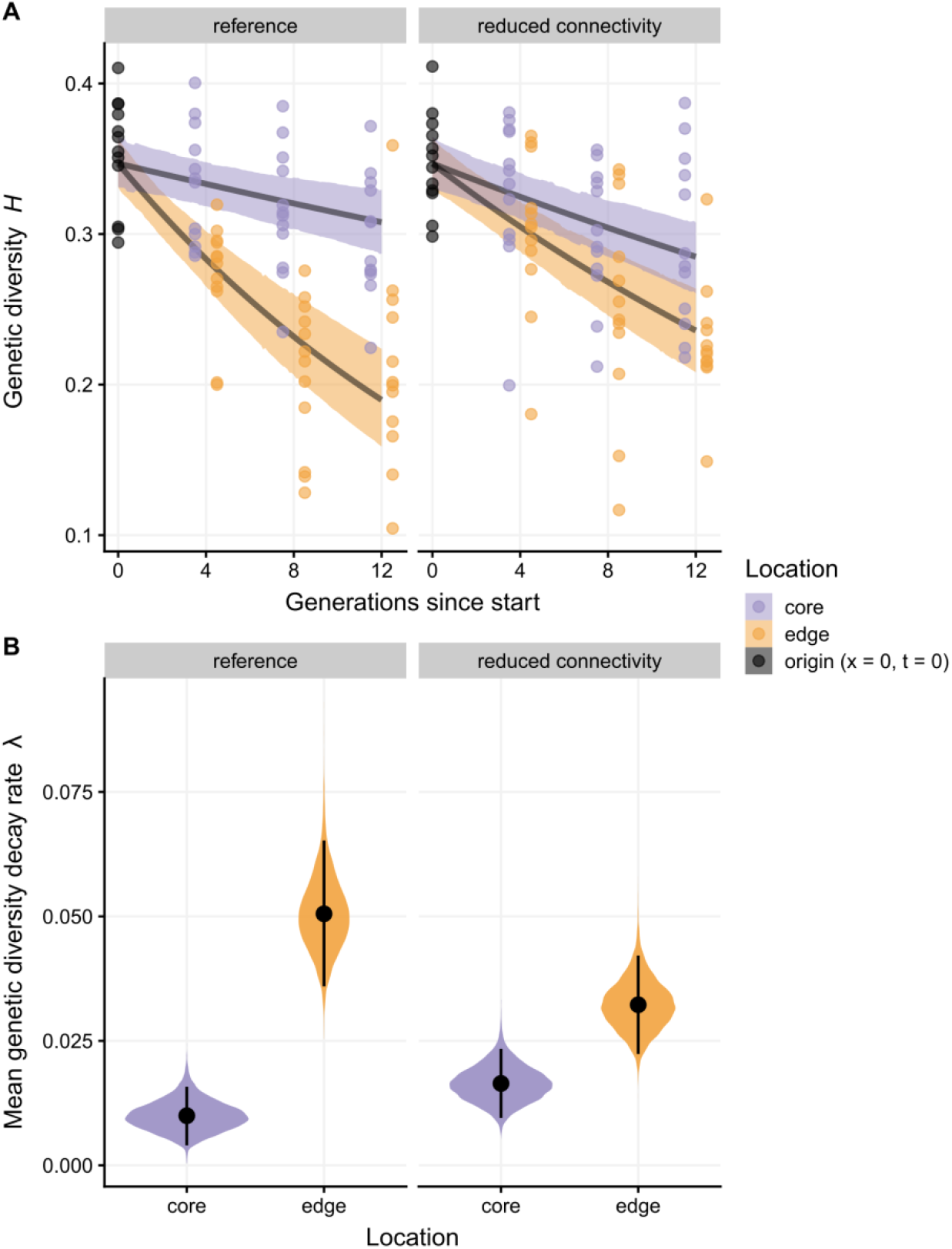
(A) Observed (points) and predicted (lines and bands) genetic diversity as a function of generations since release and landscape type. For predictions, lines are the posterior means and bands correspond to the 95% credible intervals. (B) Posterior distribution of the average genetic diversity decay rate *λ* as a function of landscape type and patch location (either the original release site (core) or the most advanced patch at the time of measure (edge)). Dots are posterior means, vertical bars 95% credible intervals.

## Discussion

By combining an individual-based model and replicated range expansions in experimental landscapes, we showed that reducing connectivity can lead to more pushed expansions. Discrepancies between the simulation and experimental results, as well as comparisons between the different simulation treatments, potentially shed light on the mechanisms at play in each context.

### Expansion velocity as a quantitative and qualitative indicator of pushed expansions

First, our simulation results largely confirm previous theoretical and empirical results regarding the effects of density-dependent dispersal and Allee effects (Birzu et al., 2018, 2019; Gandhi et al., 2019, 2016; Haond et al., 2018; Roques, Garnier, Hamel, & Klein, 2012). We show that both mechanisms lead to expansions that can be classified as “pushed” based on the velocity ratio 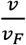 (**Fig. 2**), i.e. advancing faster than expected based on dispersal and growth at low densities. We also show that in the reference scenario, *v* can be lower than *v*_*F*_, which is expected as a result of demographic stochasticity (Brunet & Derrida, 1997; Hallatschek & Korolev, 2009), although the difference is in our case minimal (Haond et al., 2018).

In simulations, reducing connectivity led to slower expansions (lower absolute speed *v*) that were also pushed (higher velocity ratio 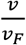) (**Fig. 4**), confirming our main prediction. Although it was not presented in connectivity terms, a similar result was recently found using a model inspired by phage-bacteria interactions (Hunter, Liu, Möbius, & Fusco, 2020), where reduced connectivity was *de facto* achieved using increased densities of phage-resistant bacteria as a barrier. In our simulations, these pushed expansions are characterized by a mean dispersal rate independent of population density; this means they would likely converge to a pulled wave as *K* tends to infinity and the effects of discreteness and stochasticity become negligible. As such, they should be considered a type of “weakly” pushed expansions (sensu Panja, 2004; Panja & van Saarloos, 2002). Nonetheless, they still advance faster than one would expect if they were pulled; further work is needed to understand the consequences of ignoring this when attempting to forecast biological invasions or climate-induced range shifts in fragmented environments.

We also confirm that, over the range of *K* we studied, increasing the equilibrium population size *K* leads to faster range expansions, as previously shown by Haond et al. (2018) for pushed expansions. Interestingly, this *K*-*v* relationship is seen in all simulated treatments, even the reference one. This may indicate that reference dispersal rates are already low enough to cause weakly pushed expansions. More likely, given the 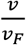 ratio of reference expansions is ≈1 or even <1 (**Fig. 4**), this may indicate that, in their case, the *K*-*v* relationship stems here from demographic (Hallatschek & Korolev, 2009), rather than dispersal stochasticity.

In experimental landscapes, contrary to simulations, expansions advanced just as fast when connectivity was reduced (**Fig. 5**). Importantly, it does not mean that reducing connectivity does not lead to more pushed expansions, on the contrary. What differentiates between pushed and pulled expansions is not absolute velocity *v* (which can be either higher or lower in pushed expansions, **Fig. 4**; Gandhi et al., 2019), but the ratio to the “equivalent” pulled wave’s velocity, so 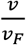. While we do not have direct quantitative information on the *v*_*F*_ of our experimental landscapes, we have some partial and indirect qualitative information. Indeed, a reduction of connectivity and thus dispersal implies by definition a reduction of *v*_*F*_, growth being equal (see also **Supplementary Material 3**). If *v*_[reference]_ ≃ *v*_[reduced]_ (**Fig. 5**) and *v*_*F*[reference]_ > *v*_*F*[reduced]_, then it follows that 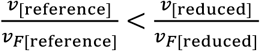. This would mean that “reduced connectivity” expansions are indeed more pushed than reference expansions in our experiment. We acknowledge that this conclusion hinges on growth rates being similar between connectivity treatments; they should be, as wasps came from the same stock populations, but evolution during expansions cannot be excluded *a priori* (Van Petegem et al., 2018). Our interpretation that reduced connectivity leads to pushed expansions in experiments is further supported by the presence of a *K*-*v* correlation that is absent in the “reference” treatment (**Fig. 6**)(Haond et al., 2018; but see simulation results above). More studies are needed to confirm these results as both our velocities and our correlations are estimated with non-negligible uncertainty, due to limited sample sizes. Nonetheless, our data show that it is possible, in principle, to make qualitative assessments of expansion “pushiness” relative to a reference, even without knowledge of the underlying dispersal or growth functions. This is very interesting for the study of biological invasions and range shifts in the wild, especially in (larger) organisms that are not easily reared and maintained in laboratory conditions. More work is needed to determine whether more quantitative insights can also be obtained from incomplete life-history and population dynamics data typical of “natural” study systems.

### Mismatches to genetic diversity predictions show pushed expansions are diverse

A key theoretical prediction is that pushed expansions lose genetic diversity at the edge more slowly than comparable pulled expansions (Birzu et al., 2019; Roques et al., 2012). While our simulations with density-dependent dispersal or Allee effects conform to this prediction (**Fig. 7**), this is not the case for the pushed expansions we generated by reducing connectivity alone: these expansions lost genetic diversity slightly faster than the pulled reference (**Fig. 7**) despite being clearly “pushed” based on velocity ratios. On the one hand, this is unsurprising, as we should expect a negative relationship between genetic diversity at expansion edges and landscape fragmentation (e.g. Hill, Hughes, Dytham, & Searle, 2006; Mona, Ray, Arenas, & Excoffier, 2014); less connectivity means new populations are founded by fewer individuals, leading to stronger founder effects and drift. On the other hand, this creates an apparent conflict between the genetic and velocity-based definitions of pushed expansions, which were implied to be intrinsically correlated (e.g. Birzu et al., 2018). We suggest that, although “weakly” pushed expansions can behave like “classical” pushed expansions with respect to speed, they are actually qualitatively different, and we should not expect all theoretical predictions based on “classical” pushed expansions to apply to them. Further research is thus needed to better pinpoint where “classical” and “weakly” pushed expansions diverge or converge in their dynamics. Setups using natural inter-individual variation in dispersal (Saastamoinen et al., 2018; Schreiber & Beckman, 2020; Stevens, Pavoine, & Baguette, 2010) to decouple the effects of reduced connectivity itself from those of increased dispersal variability might be particularly useful here.

In experimental landscapes, reducing connectivity did slow the decay of genetic diversity at the edge of the expansion (**Fig. 8**). Given population genetics models predict a faster loss of diversity with increased fragmentation when expansions are not pushed (Mona et al., 2014), this result is strong evidence that our experiments did generate (more) pushed expansions, this time in the “classical” sense. On the other hand, experiments are here in direct contradiction with our simulation results. This mismatch may indicate increased dispersal stochasticity is not the only mechanism at play here.

Based on simulation results (**Fig. 7**), we must first consider the possibility that experimental landscapes with reduced connectivity lose genetic diversity more slowly simply because they exhibit larger core populations, rather than due to a shift in regime from (more) pulled to (more) pushed expansions. There is however no clear evidence that changes in connectivity led to changes in equilibrium population sizes (see Results), so we do not believe this is the main driver of our genetic diversity results. In addition, even if there were evidence of changes in *K*, this would have led to differences in genetic diversity not only in edge but also in core populations, which we did not observe (**Fig. 8**).

From a dispersal ecology standpoint, one of the most obvious differences between our simulations and experiments is that we did not allow simulated individuals to adjust their dispersal-density decision rules as a function of their environment. Dispersal decisions are multicausal, with many factors interacting to shape the benefits-cost balance of dispersal (Matthysen, 2012). For instance, emigration by *Notonecta undulata* backswimmers is density-dependent in parasite-free ponds, but density-independent in mite-infected ponds (Baines, Diab, & McCauley, 2020). In *Erigone atra* spiders, dispersal by rappelling depends on the interaction of sex, density and local sex-ratios (De Meester & Bonte, 2010). In *Tribolium castaneum* beetles, dispersal is less density-dependent out of lower-quality habitat (Endriss et al., 2019; Van Allen & Bhavsar, 2014). In our context, one might wonder how reduced connectivity and density-dependent dispersal might interact. Theory predicts that density-dependent emigration should be more advantageous when connectivity is reduced (more precisely, when costs of dispersal are higher; Travis, Murrell, & Dytham, 1999; Rodrigues & Johnstone, 2014). In line with these predictions, dispersal was density-dependent in *Tribolium castaneum* metapopulations with low connectivity, density-independent in metapopulations with high connectivity (Govindan, Feng, DeWoody, & Swihart, 2015). If we want to see whether simulations and experiments actually agree, these previous studies suggest we should actually compare our experimental treatments to the “reference” vs “reduced connectivity + density-dependent dispersal” pair of treatments in our simulations, rather than the “reference” vs “low connectivity alone” pair. Confirming this hypothesis, the “reduced connectivity + density-dependent dispersal” simulated expansions are a better match to the experiments in that (i) while pushed, their absolute speed *v* is closer to reference expansions, and (ii) they do lose genetic diversity slower than reference expansions. For this mechanism to plausibly apply to our experimental system, *Trichogramma* wasps need to be able to detect when density and competition increase locally, and adjust their spatial behavior in response, which evidence suggests they do (Wajnberg, Fauvergue, & Pons, 2000). Importantly, the hypothesis that low connectivity drives changes in the density-dependence of dispersal itself, leading to “true” and not simply “weakly” pushed expansions, does not imply these changes result from evolution during the expansion. Indeed, in at least four out of the five experiments cited above, the context-specific adjustments to the dispersal-density function involved plastic responses (Baines et al., 2020; De Meester & Bonte, 2010; Endriss et al., 2019; Van Allen & Bhavsar, 2014). Further experiments under standardized densities and connectivity are needed to determine whether our results are explained by plastic adjustments or evolutionary changes.

## Conclusion

In this study, we demonstrated a new mechanism for generating pushed(-like) expansions (**Fig. 1**). We showed that the genetic and velocity aspects of the pushed/pulled distinction, which have been assumed to be tightly linked (Birzu et al., 2018) can be decoupled when (weakly) pushed expansions are caused solely by reduced connectivity. We also confirm pushed expansions can be detected and analyzed in systems with (relatively) low population sizes, even though theory has mostly been developed on much higher population sizes. This means the pushed-pulled framework may be valuable to help understand and better predict range expansions and shifts in a broad range of taxa and contexts (including under natural conditions). As ecologists and evolutionary biologists are becoming more aware and more interested in the potential effects of pushed expansions (Miller et al., 2020; Williams, Hufbauer, & Miller, 2019), our results suggest we must be careful about which questions we ask, and whether the dimensions of “pushiness” we collect data on are the appropriate ones. For instance, assuming an expansion loses less genetic diversity simply because its measured velocity ratio is higher may not always be appropriate (**Figs 4, 7**). This increased focus on ecological pushed expansions is nonetheless welcome, as our results highlight more studies are needed to better understand how their impacts depend on the underlying causal mechanisms.

Our experiment shows that, in some cases, reducing connectivity may limit the loss of genetic diversity without impeding spread rates. Founder effects and low genetic diversity are associated with lower adaptive potential and lower chances of population persistence (e.g. Szűcs, Melbourne, Tuff, Weiss-Lehman, & Hufbauer, 2017). Our results thus lead to the somewhat counterintuitive conclusion that reducing connectivity might, in some cases, help expanding species. This conclusion has strong implications for the management of invasive species and the conservation of species undergoing climate-induced range shifts, as one of the main causes of pushed expansions, positive density-dependent dispersal, is frequent in nature (Harman et al., 2020; Matthysen, 2005). Finally, density-dependent dispersal (Fronhofer, Gut, & Altermatt, 2017; Travis, Mustin, Benton, & Dytham, 2009) and Allee effects (Datta, Korolev, Cvijovic, Dudley, & Gore, 2013; Erm & Phillips, 2020) themselves may evolve during range expansions, and the effects of habitat fragmentation on dispersal ecology and evolution are abundantly documented (Cote et al., 2017; Jacob, Laurent, Morel-Journel, & Schtickzelle, 2020). For these reasons, we call for more systematic eco-evolutionary studies of context-dependent dynamics during range expansions and shifts.

## Data accessibility

Data, Netlogo model code and R scripts to reproduce all analyses presented in this manuscript are available on GitHub (https://github.com/mdahirel/pushed-pulled-2020-dynamics, release v2.0) and archived in Zenodo (https://doi.org/10.5281/zenodo.3969988). Other Supplementary Materials are available from the same sources.

## Acknowledgements

We thank participants to the 2019 conference of the British Ecological Society for their helpful comments, as well as Géraldine Groussier for letting us use her picture of *Trichogramma brassicae*. Version 4 of this preprint has been peer-reviewed and recommended by Peer Community In Evolutionary Biology (https://doi.org/10.24072/pci.evolbiol.100118).

