## Supplementary material for "Shifts from pulled to pushed range expansions caused by reduction of landscape connectivity": supplementary_material.html


### Supplementary Material for “Shifts from pulled to pushed range expansions caused by reduction of landscape connectivity”

###### Maxime Dahirel, Aline Bertin, Marjorie Haond, Aurélie Blin, Eric Lombaert, Vincent Calcagno, Simon Fellous, Ludovic Mailleret, Thibaut Malausa, Elodie Vercken (code by M. Dahirel)

**Note:**  The full code for the following document and the analyses it describes, as well as for all analyses described in the main text, is available on GitHub here and Zenodo: .

### S1 - Creation of genetic lines

*Trichogramma brassicae* wasps used in the experiment were derived from hybrid populations created in 2013 by mixing parental strains sampled in 24 sites in Western Europe. These parental strains were initially collected for a project on artificial selection and phenotype amelioration of *T. brassicae* strains for biological control. Isoline populations were then maintained under standard conditions between 2014 and 2016 (18 ± 0.5 °C, 65 ± 10% relative humidity, L:D 18:6), for approximately 20 generations.

To generate independent experimental replicates with comparable levels of initial genetic diversity, and to allow for evolutionary dynamics, we created three genetic mixes from three isoline populations each (using nine distinct isoline populations in total). We followed the protocol from Fellous *et al.* (2014) to create mixes with homogeneous genetic background (i.e. similar representation of each of the three ancestral isolines). First, we performed all possible two-by-two crosses between the three isolines. The F1 progeny of each two-by-two cross was then crossed with the third remaining isoline. The F2 individuals from these crosses were then inbred for 2 generations to favour recombination. Individuals from these F4 were then mixed all together in equal proportions to form each final genetic mix. While they did differ from each other, all three mixes had broadly similar levels of genetic diversity at the start of the experiments (**Supplementary Figure S1.1**).

**Supplementary Figure S1.1** Genetic diversity (multilocus expected heterozygosity) as a function of genetic mix at the start of the experiment. One dot = one experimental landscape. The y axis is scaled to show the full range of genetic diversities observed during the course of the experiment, for reference (see main text)

### S2 - Validating computer estimates of *Trichogramma* population sizes

For analyses asking for population sizes, the proportion of parasitized eggs was estimated based on egg strip photographs using ImageJ (Abramoff, Magalhães, & Ram, 2004) and the `CODICOUNT` plugin (Perez, Burte, Baron, & Calcagno, 2017). `CODICOUNT` estimates the number of pixels assigned to parasitized host eggs, unparasitized host eggs and background after training. We tested each photograph using 4 macros, each trained on a different set of pictures (from generations 1 2, 3 and 13 respectively) and each potentially having its own systematic bias, combined to better approximate real rates. Proportions of pixels identified as “egg” that were also identified as “parasitized” were used as our estimate of parasitism rate/ population size.

To assess the validity of these automated counts, we manually counted parasitized egg plates from 197 patches (out of 2725 photographed for the experiment) roughly spanning the full range of possible population sizes (human count range: 17 to 443; number of hosts per population ≈ 450).

We then used a bivariate generalized linear mixed model approach to compare computer estimates and human counts, with a Beta family used for the former and a binomial distribution for the latter. Both models contained a fixed-effect intercept as well as random intercepts of population/patch identity, which were correlated. This approach accounts for the data structure and the fact both human- and computer-derived counts are likely estimated with error (Perez et al., 2017). It allows us to estimate the correlation between human counts and “consensus” computer counts obtained from pooling information from all macros.

Estimated and human-observed parasitism rates were highly correlated both on the latent logit scale (*r*= 0.87 [0.83, 0.9]) and on the observed scale (*r* = 0.9 [0.89, 0.91]). There was also no evidence of bias (predicted mean difference between “human” and “computer” parasitism rates, in % of hosts per patch = 0.34% [-0.22, 0.88])

We then refitted this model, this time adding a fixed effect of the macro in the “computer estimates” submodel, to determine whether macros presented systematic biases. We found that the overall absence of bias of the “consensus” estimates (see above) actually resulted from the majority of macros slightly underpredicting human-observed parasitism rates while one consistently overpredicted them (**Supplementary Figure S2.1**).

**Supplementary Figure S2.1** Posterior mean estimated parasitism rate, by computer macro, for the 197 plates used to compare human and computer counts. Horizontal lines denote the mean (bold line) and 95% credible interval (dotted lines) for the posterior mean parasitism rate based on human counts.

### S3 – Validation of the experimental design

To determine whether our experimental reduction of structural connectivity did limit individual movement (i.e. functional connectivity), we used data from the first generation of testing, i.e. the only time the patch of origin of all parents can be known with certainty.

We used parasitism rates to calculate for each landscape × macro the average egg location (in patches from release) and used it as a proxy of dispersal distance (works because dispersal can be define as more or less parent-offspring distance). We analysed this distance using a Beta generalised linear mixed model (dividing it by 4, the maximal possible distance, to ensure it stayed between 0 and 1). The model included a fixed effect of treatment, as well as a random effect of macro (random intercept) to account for uncertainty in our estimates of parasitism and distances, more specifically for macro-level systematic biases.

As expected, we found that released wasps did lay their eggs closer from release sites in landscapes with reduced connectivity compared to reference landscapes (**Supplementary Figure S3.1**).

**Supplementary Figure S3.1.** Left: Average egg-laying distance from release patch as a function of treatment. Colored areas: posterior distributions of the averages; black dots and segments: posterior mean and 95% credible intervals; empty colored dots: estimated values for each macro; full colored dots: averages across macros for each replicate. Right: posterior difference between the two experimental treatments.

### S4 - Microsatellite genotyping, detailed methods

#### S4.1 - Design of the microsatellite markers and multiplex PCR assay

Microsatellite DNA libraries were produced following the method described in Malausa *et al.* (2011). This method consists of (i) the isolation of DNA fragments containing microsatellite motifs by multiplex microsatellite enrichment (obtained via hybridisation to biotin-labelled oligonucleotides with the motifs (AG)_10_, (AC)_10_, (AAC)_8_, (AGG)_8_, (ACG)_8_, (AAG)_8_, (ACAT)_6_ and (ATCT)_6_) and (ii) the pyrosequencing of the fragments (454 GS-FLX Titanium). This protocol was applied to a sample containing DNA extracted from about 100 *Trichogramma brassicae* individuals.

The obtained DNA dataset was analyzed and primers for microsatellite amplification were designed using the QDD program (Meglécz et al., 2010), version 2.1. The default QDD parameters were used, except for: (i) the minimum percentage similarity of sequences used for the construction of consensus sequences was 90%, (ii) the proportion of sequences that must have the same base on the aligned site to accept it as a consensus was 0.66, (iii) the minimum length of PCR product for primer design was 80bp, (iv) the maximum length of PCR product for primer design was 500, (v) the maximum length of a primer was 32, (vi) the maximum acceptable difference between the melting temperatures of primers was 4°C.

Nineteen microsatellite markers were developed by testing and multiplexing part of the PCR primers selected after the QDD analysis. The main criteria for microsatellites sequences selection and primers design used were (i) the motif of the microsatellite they target, to obtain a balanced frequency of motifs, (ii) microsatellites with at least eight repeats, (iii) the size of the expected PCR products (we selected primers amplifying markers homogeneously distributed in terms of size within the range 80 bp - 500 bp).

#### S4.2 - Genotyping and microsatellites description

All sequenced *T. brassicae* wasps were stored in 70% ethanol before DNA extraction. DNA was extracted from each individual wasp, either with the prepGem Insect DNA Extraction kit from Zygem (extraction volume of 15µL per individual, using 0.375µL of prepGem enzyme; samples were placed in a thermocycler for 2 hours at 75°C and 5 minutes at 95°C), or with the Quick Extract DNA Extraction Solution from Lucigen (extraction volume of 15µL per individual; samples were placed in a thermocycler for 10 minutes at 65°C and 2 minutes at 98°C). All DNA extracts were stored at -20°C.

For microsatellite amplification, 2µl of genomic DNA were used within a 10µL final PCR reaction consisting of 5µL of the QIAGEN PCR multiplex solution, 1µL of the primer mix of the 19 primers reverse and forward (1µM each), and water. PCR conditions were as followed: initial denaturation at 95°C for 15 min; 30 cycles of denaturation (94°C, 30s), annealing (60°C, 90s) and elongation (72°C, 60s); final extension at 60°C for 30 min. For each individual, 1.5µL of the PCR product was mixed with 9.8µL of a mix obtained from 1mL HiDi formamide and 20µL of 500 LIZ Size Standard (Applied Biosystems). Products were then electrophoresed using an ABI PRISM 3130XL sequencer (Applied Biosystems). Finally, genotypes were scored using Gene Marker version 1.75 (SoftGenetics).

All loci are polymorphic when looking at the entire dataset (**Supplementary Table S4.1**). Information on each marker, including allelic richness over the whole set of genotyped individuals, is presented **Supplementary Table S4.1**. Information about genetic diversity and whether alleles frequencies followed the Hardy-Weinberg equilibrium **at the start of the experiment** is presented Supplementary Table S4.2.

Summary data **Table S4.2** confirm that all 3 source strains (genetic mixes) are polymorphic and diverse at the majority of the loci chosen, and that only a limited number of loci are polymorphic in 1 strain and fixed in the other 2. One locus has H0(t = 0) = 0 for all strains. However, we note that all loci do present variation in later generation samples (a prerequisite for keeping the locus for the analysis; data not shown here). So, barring multiple mutations, this seemingly fixed locus must simply represent sampling error at t = 0.

**Supplementary Table S4.1**: Primer pairs used for the PCR set of the multiplex panel. For each of the 19 loci, the table includes: the repeat motifs in the sequence used to design primers, sequence and fluorescent dye label used for each primer, the observed number of alleles and the allelic size range.

|  | | Primer sequences (5’—3’) | |  | | |
| --- | --- | --- | --- | --- | --- | --- |
| Locus | Repeat motif | Forward | Reverse | Fluorescent dye | number of alleles | Size range (bp) |
| TB17 | (AACG)_6_ | CCTCAAAGCCGACGTGTAG | ACATAAGCTTCGGTGGTTGC | FAM | 4 | 109 - 123 |
| TB45 | (AAC)_10_ | GAGGCTTCAATTGGTTTTCG | CTTACGGTATCGTGCAGCAG |  | 2 | 193 - 196 |
| TB73 | (AGC)_10_ | CAGACACAGACACGCACCTT | TCTCATTGTGTGATTTATGCGA |  | 5 | 265 - 285 |
| TB84 | (AG)_12_ | CAGCAGCGACGAGTACAAAG | TATAACGATTCGGAGTCGGC |  | 2 | 299 - 301 |
| TB100 | (AC)_14_ | CGAAGCTGCTCTCGAATAGG | GGGAAAGTGAACGAGCGATA |  | 7 | 350 - 362 |
| TB112 | (AAC)_15_ | AATTCCTAGTCGGAAGCCG | AGTGTACACGCATACACGGG |  | 2 | 382 - 385 |
| TB5 | (AGC)_11_ | TCGAAGTCGAAGCTCGTTG | GACGTCGAAACTTGATGCTG | VIC | 9 | 73 - 89 |
| TB10 | (AAG)_8_ | GGCCTATGCGTATCTTGGAC | TGCTACCGTCGTTGAACTTG |  | 4 | 100 - 111 |
| TB39 | (AG)_13_ | ATATACCAGCGGAGCAAAGC | GCGTGCATAAAACAGTGGTC |  | 4 | 175 - 179 |
| TB107 | (AC)_14_ | AACGAGGCTACACGTGGATAA | CACAAGTTGCACGAGTAACGA |  | 4 | 363 - 373 |
| TB3 | (AG)_10_ | AACTTCTGTGCAAAGCCTCG | CAAGAGAGCGAGCGAGAGAG | NED | 5 | 81 - 92 |
| TB15 | (AG)_19_ | TGATAGAGAAGGGCAAACGG | ACGGCAGTCTGCAAGAGAAG |  | 6 | 121 - 139 |
| TB48 | (ACT)_10_ | TATACGCCGACGAGTTTGG | TGCTGCAGGTCAGGTGTCTA |  | 3 | 182 - 201 |
| TB75 | (AG)_20_ | GCGCATACCTAGCACCAAGT | CATACCGTACGGCAACGAC |  | 9 | 246 - 279 |
| TB101 | (AAC)_14_ | TATGTACGCAGCTACACCGC | CTCGGCCATTCTTTATTTCG |  | 10 | 320 - 367 |
| TB24 | (ACT)_8_ | TTCTTTCACGCTCACACCC | GTTTTGCAGCTCGTGACAAC | PET | 3 | 133 - 148 |
| TB42 | (AC)_12_ | CTTCTAGCTTGCCTCCATCG | GAGCGGAGCATTTCAATCTT |  | 4 | 178 - 194 |
| TB76 | (ACT)_13_ | TACATATGCAGGCGATACACTGA | GTGCACAGGCTGCTTCATATAC |  | 3 | 275 - 291 |
| TB99 | (ACT)_15_ | GTAGATTGCAGAAGCGGTGC | GTCCGATCGATCCAATGGT |  | 6 | 334 - 363 |

**Supplementary Table S4.2**: Population genetics statistics **at the start of the experiment** (Generation 0 samples). At each locus described **Supplementary Table S4.1**, the number of individuals successfully genotyped (*N*, excluding missing alleles),the observed heterozygosity (*H_O_*), the expected heterozygosity (*H_E_*, Nei (1973)) and the *p*-value for the exact test of Hardy–Weinberg equilibrium were computed separately for each of the 3 genetic mixes used (Mix1, Mix2, Mix3; See Supplementary Material S1). Statistics were computed using the `adegenet` (Jombart, 2008) and `pegas` (Paradis, 2010) R packages. Note that all loci were found to be polymorphic when analysing the whole set of wasps used in the experiment (See **Supplementary Table S4.1**).

| Locus | *N* | *H_O_* | *H_E_* | HWE *p*-value |
| --- | --- | --- | --- | --- |
| TB10 | 100, 118, 80 | 0.83, 1, 1 | 0.58, 0.5, 0.59 | 0, 0, 0 |
| TB100 | 132, 120, 127 | 0.49, 0.47, 0.45 | 0.49, 0.41, 0.48 | 0.54, 0.17, 0.69 |
| TB101 | 131, 119, 127 | 0.04, 0, 0 | 0.59, 0, 0.44 | 0, –, 0 |
| TB107 | 131, 119, 127 | 0.42, 0.43, 0.18 | 0.41, 0.44, 0.16 | 0.82, 0.66, 0.61 |
| TB112 | 132, 120, 127 | 0, 0.44, 0.52 | 0, 0.48, 0.5 | –, 0.48, 0.74 |
| TB15 | 132, 119, 127 | 0.45, 0.39, 0.63 | 0.45, 0.45, 0.65 | 1, 0, 0.97 |
| TB17 | 132, 119, 127 | 0, 0.61, 0 | 0, 0.48, 0 | –, 0.01, – |
| TB24 | 132, 119, 127 | 0.48, 0, 0 | 0.5, 0, 0 | 0.6, –, – |
| TB3 | 132, 119, 127 | 0.49, 0.55, 0.57 | 0.45, 0.45, 0.62 | 0.3, 0.04, 0.09 |
| TB39 | 132, 120, 127 | 0, 0.44, 0.51 | 0, 0.47, 0.53 | –, 0.46, 0.16 |
| TB42 | 132, 120, 127 | 0.54, 0.49, 0.31 | 0.53, 0.54, 0.31 | 0.88, 0.28, 0.38 |
| TB45 | 132, 120, 127 | 0.35, 0.12, 0.46 | 0.34, 0.19, 0.5 | 1, 0, 0.49 |
| TB48 | 132, 120, 127 | 0, 0, 0 | 0, 0, 0 | –, –, – |
| TB5 | 129, 119, 112 | 0.4, 0.13, 0.02 | 0.58, 0.29, 0.54 | 0, 0, 0 |
| TB73 | 132, 120, 127 | 0.5, 0.5, 0.61 | 0.5, 0.46, 0.66 | 1, 0.42, 0.04 |
| TB75 | 132, 120, 127 | 0.48, 0.36, 0.5 | 0.46, 0.4, 0.5 | 0.85, 0.24, 0.86 |
| TB76 | 132, 120, 127 | 0.46, 0.43, 0.63 | 0.45, 0.43, 0.64 | 0.89, 1, 0.92 |
| TB84 | 132, 120, 127 | 0.45, 0, 0 | 0.42, 0, 0 | 0.54, –, – |
| TB99 | 132, 120, 127 | 0.41, 0.44, 0.59 | 0.44, 0.48, 0.62 | 0.41, 0.38, 0.36 |

### S5 - Detailed description of models

We here describe the full structure of the models presented in the main text and supplementary materials, as well as associated prior choices. We try to follow throughout the same notation conventions as McElreath (2020). Unless noted, variable names and indices are reset from one model to the next, for simplicity, but similar names always refer to similar types of variables (fixed effects/random effects/correlations…). All Beta models descriptions use the (mean, precision) parametrisation of the Beta distribution. We use the \(\mathrm{Half\mbox{-}Normal}(0,\sigma)\) notation to denote a half-normal distribution based on a \(\mathrm{Normal}(0,\sigma)\) distribution.

#### S5.1 - Genetic diversity, experimental data

We can describe the genetic diversity (expected heterozygosity) \(H\_{x,i,t}\) of a patch in replicate \(i\), with \(x\) denoting the location of that patch (0 for core patches, 1 for edge/front patches) and \(t\) the number of generations since release, the following way:

\[
\mathrm{logit}(H\_{x,i,t}) \sim \mathrm{StudentT}(\mathrm{logit}(\mu\_{x,i,t}) , \sigma\_{d} , \nu ),
\] where \(\mu\_{x,i,t}\) is the mean genetic diversity, \(\sigma\_{d}\) the distribution scale parameter and \(\nu\) the number of degrees of freedom.

Based on theory, we expect average genetic diversity to decline exponentially with time:

\[
\mu\_{x,i,t} = \mu\_{i,t=0} \times \exp(-\lambda\_{x,i} \times t).
\]

The two parameters \(\mu\_{i,t=0}\) (initial diversity) and \(\lambda\_{x,i}\) (rate of decline) depend on replicate, treatment (reference or reduced connectivity) and location (core \(x = 0\) or edge \(x = 1\)) as follows:

\[
\mathrm{logit}(\mu\_{i,t=0}) = \beta\_{0} + \alpha\_{i}, \\
\log\_{10}(\lambda\_{x,i}) = \beta\_{1[\text{TREATMENT}[i]]}+\beta\_{2[\text{TREATMENT[i]}]}(x-0.5) + \gamma\_{i} + \zeta\_{i}(x-0.5)
\] (with \(\mathrm{logit}\) and \(\log\_{10}\) transformations here to ensure both parameters stay within bounds).

Initial diversity is thus independent of treatment and location, and only depends on replicate identity through a random effect \(\alpha\_{i}\), while the rate of decline \(\lambda\) depends on treatment, location and their interaction. (Note the centring of \(x\) in the formula, and that \(x\) is set to 0.5 at \(t\) = 0, so that the effect of location on \(\lambda\) cancels out at the start of the experiment, when “core” and “edge” are not distinct). In addition, both the average decline rate and the edge-core difference depend on replicate, which is accounted for through the random effect parameters \(\gamma\_{i}\) and \(\zeta\_{i}\) respectively.

The random effects are distributed as follows:

\[
\begin{bmatrix}
\alpha\_{i} \\
\gamma\_{i} \\
\zeta\_{i} \\
\end{bmatrix} \sim
\mathrm{MVNormal}(
\begin{bmatrix}
0\\
0\\
0\\
\end{bmatrix},\boldsymbol{\Omega}),
\] with \(\boldsymbol{\Omega}\) the covariance matrix:

\[
\boldsymbol{\Omega} =
\begin{bmatrix}
\sigma\_{\alpha} & 0 & 0 \\
0 & \sigma\_{\gamma} & 0 \\
0 & 0 & \sigma\_{\zeta} \\
\end{bmatrix}
\times \boldsymbol{\mathrm{R}} \times
\begin{bmatrix}
\sigma\_{\alpha} & 0 & 0 \\
0 & \sigma\_{\gamma} & 0 \\
0 & 0 & \sigma\_{\zeta} \\
\end{bmatrix},
\] where \(\sigma\_{\alpha}\), \(\sigma\_{\gamma}\) and \(\sigma\_{\zeta}\) are the random effect standard deviations for each parameter, and \(\boldsymbol{\mathrm{R}}\) the corresponding correlation matrix.

We used weakly informative priors mostly following McElreath (2020). For the fixed intercept of the initial diversity \(\beta\_{0}\), which corresponds to the logit of a proportion, we used a \(\mathrm{Normal}(0,1.5)\), which gives a relatively flat prior when converted back to proportions. For all other fixed effects parameters \(\beta\_{j|j > 0}\), we used a \(\mathrm{Normal}(0, 1)\) prior (note that results are overall insensitive to fixing both standard deviations to 1 or to 1.5). All standard deviation parameters \(\sigma\) (including the distributional standard deviation \(\sigma\_{d}\)) were attributed the same \(\mathrm{Half\mbox{-}Normal}(0,1)\) prior. For the random effect correlation matrix \(\boldsymbol{\mathrm{R}}\) we used a \(\mathrm{LKJCorr}(2)\) prior, while we used a \(\mathrm{Gamma}(2,0.1)\) for the degrees of freedom \(\nu\), based on Stan developers recommendations (https://github.com/stan-dev/stan/wiki/Prior-Choice-Recommendations).

An alternative model can be written, where \(H\_{x,i,t} \sim \mathrm{Beta}(\mu\_{x,i,t}, \phi)\), but it has a lower predictive performance (see Methods in the main text, and details in linked code (main script 1)). Following recommendations for weakly informative priors (https://github.com/stan-dev/stan/wiki/Prior-Choice-Recommendations), we use there a \(\mathrm{Half\mbox{-}Normal}(0,1)\) prior on \(1/\phi\) (all other parameters and priors are as described above).

#### S5.2 - Genetic diversity, simulated data

In our simulations, it is not possible to analyse genetic diversity using the model(s) described in S5.1, in large part because simulated diversity data contains many zeroes, which means models using both logit transformations and Beta distributions are inappropriate.

As an alternative, we can use the model proposed by Gandhi *et al.* (2019) in which, for a given treatment and location combination \(j\) at a given time \(t\) (in generations from release), the expected among-replicate variance in the frequencies of either one of the two alleles \(V\_{j,t}\) can be described this way:

\[ V\_{j,t} \sim \mathrm{Beta}(\mu\_{j,t}, \phi),\]

\[ \mu\_{j,t} = V\_{\max} \times (1 - \exp(-\lambda\_{j} \times t)),\]

where \(V\_{max}\) is both the asymptotic variance (once all patches have reached fixation) and the product of initial allele frequencies (and so only depends on the initial allelic distribution, not on treatment or patch type), and \(\lambda\) a rate of decay of genetic diversity, which varies between treatments and locations (core vs. edge) as in the “experimental” model.

We put the same priors on \(\log\_{10}(\lambda)\) and \(1/\phi\) as in the experimental models (\(\mathrm{Normal}(0, 1)\) and \(\mathrm{Half\mbox{-}Normal}(0, 1)\), respectively), for the same reasons. We put a stronger, informative prior on \(\mathrm{logit}(V\_{max})\) :

\[ \mathrm{logit}(V\_{max}) \sim \mathrm{Normal}(\mathrm{logit}(0.25), 0.5), \] as the expected asymptotic among-replicate variance is 0.25 for the case of a randomly distributed and randomly fixating locus with two alleles (\(p(1-p)\) with \(p\) = 0.5).

#### S5.3 - Front velocity, experimental and simulated data

In both experimental and simulation data, the location \(X\_{i,t}\) of the expansion front of the \(i\)th replicate after \(t\) generations can be modelled as follows:

\[ X\_{i,t} \sim \mathrm{Log\mbox{-}Normal}(\log (v\_{i,t} \times t), \sigma\_{d}),\]

where \(v\_{i,t}\) denotes the expansion speed over the \(t\) generations since release, and \(\sigma\_{d}\) is the residual standard deviation of the log-distances.

The expansion speed \(v\_{i,t}\) can then be modelled as follows:

\[v\_{i,t} = v\_{i,t \to \infty} + (v\_{i,t=1} - v\_{i,t \to \infty}) \times \exp(-\lambda\_{i} \times (t -1)),\]

that is, moving from an initial speed \(v\_{i,t=1}\) to an equilibrium speed \(v\_{i,t \to \infty}\), with the rate of exponential “decay” to the asymptotic speed being \(\lambda\_{i}\).

In the experiment, all three parameters depend on treatment, and on replicate identity:

\[
\log(v\_{i,t=1}) = \log(v\_{\text{TREATMENT}[i], t=1}) + \alpha\_{i}, \\
\log(v\_{i,t \to \infty}) = \log(v\_{\text{TREATMENT}[i], t \to \infty}) + \gamma\_{i},\\
\log\_{10}(\lambda\_{i}) = \log\_{10}(\lambda\_{\text{TREATMENT}[i]}) + \zeta\_{i}.
\]

In simulated data, we fixed \(v\_{i,t=1} = 1\) (see main text for rationale), we logit-transformed \(v\_{i,t \to \infty}\) rather than log-transforming it (because speeds were constrained between 0 and 1):

\[
\mathrm{logit}(v\_{i,t \to \infty}) = \mathrm{logit}(v\_{\text{TREATMENT}[i], t \to \infty}) + \gamma\_{i},
\] and we rescaled \((t -1)\) to \((t - 1) /10\) (so expressed elapsed time in 10s of generations), to facilitate convergence while keeping the same weakly informative priors. The model is otherwise unchanged.

In both simulated and experimental data, the random effects \(\alpha\_{i}\), \(\gamma\_{i}\) and \(\zeta\_{i}\) are distributed as follows:

\[
\begin{bmatrix}
\alpha\_{i} \\
\gamma\_{i} \\
\zeta\_{i} \\
\end{bmatrix} \sim
\mathrm{MVNormal}(
\begin{bmatrix}
0\\
0\\
0\\
\end{bmatrix},\boldsymbol{\Omega}),
\]

with \(\boldsymbol{\Omega}\) the covariance matrix:

\[
\boldsymbol{\Omega} =
\begin{bmatrix}
\sigma\_{\alpha} & 0 & 0 \\
0 & \sigma\_{\gamma} & 0 \\
0 & 0 & \sigma\_{\zeta} \\
\end{bmatrix}
\times \boldsymbol{\mathrm{R}} \times
\begin{bmatrix}
\sigma\_{\alpha} & 0 & 0 \\
0 & \sigma\_{\gamma} & 0 \\
0 & 0 & \sigma\_{\zeta} \\
\end{bmatrix},
\]

where \(\sigma\_{\alpha}\), \(\sigma\_{\gamma}\) and \(\sigma\_{\zeta}\) are the random effect standard deviations for each parameter, and \(\boldsymbol{\mathrm{R}}\) the corresponding correlation matrix.

We used \(\mathrm{Normal}(0,1)\) priors for all fixed effects parameters (except \(\mathrm{logit}(v\_{\text{TREATMENT}[i], t \to \infty})\) for simulated data, \(\mathrm{Normal}(0,1.5)\)). We used \(\mathrm{Half\mbox{-}Normal}(0,1)\) priors for all standard deviations \(\sigma\) and a LKJ prior \(\mathrm{LKJcorr}(2)\) for the correlation matrix \(\boldsymbol{\mathrm{R}}\)

Alternatively, one can assume (based on theory) that speed converges to the asymptote following an power law, rather than an exponential:

\[
v\_{i,t} = v\_{i,t \to \infty} + (v\_{i,t=1} - v\_{i,t \to \infty}) \times t^{-\lambda\_{i}},
\]

(with all other elements of the models remaining otherwise unchanged). However, model comparison (see code) show this performs much worse than the exponential in the context of the simulated data.

#### S5.4 - Population size in core patch, experimental data

We first assumed the parasitism rate in the release patch \(P\_{i,t,m}\) observed by macro \(m\) in replicate \(i\) after \(t\) generations can be modelled using a Beta model as follows:

\[
P\_{i,t,m} \sim \mathrm{Beta}(K\_{i,m}, \phi), \\
\mathrm{logit}(K\_{i,m}) = \beta\_{\text{TREATMENT}[i]} +\eta\_{i} + \theta\_{m},\\
\eta\_{i} \sim \mathrm{Normal}(0,\sigma\_{\eta}), \\
\theta\_{m} \sim \mathrm{Normal}(0,\sigma\_{\theta}),
\] where \(K\) are the mean carrying capacities, \(\eta\) denote random effects of replicate and \(\theta\) random effects accounting for systematic between-macro differences. Priors were the same as in previous Beta models, that is a \(\mathrm{Normal}(0,1.5)\) for \(\beta\)s, and a \(\mathrm{Half\mbox{-}Normal}(0,1)\) for both random effect standard deviations \(\sigma\) and for \(1/\phi\).

However, posterior checks revealed this model predicted more variability than was in the data, like for genetic experimental data. So, like with genetic experimental data, we shifted to a Student model on logit-transformed data:

\[
\mathrm{logit}(P\_{i,t,m}) \sim \mathrm{StudentT}(\mathrm{logit}(K\_{i,m}), \sigma\_{d} , \nu),
\] with everything else unchanged. Priors are as in the previous model, so a \(\mathrm{Normal}(0,1.5)\) for \(\beta\)s, and a \(\mathrm{Half\mbox{-}Normal}(0,1)\) for random effect standard deviations \(\sigma\). We again used a \(\mathrm{Gamma}(2,0.1)\) prior for the Student-specific \(\nu\).

#### S5.5 - Correlation between \(K\_{i}\) and \(v\_{i,t \to \infty}\), experimental data

To analyse the among-replicate correlation between the asymptotic speed \(v\_{i,t \to \infty}\) and the carrying capacity \(K\_{i}\), we simply combine the two univariate models described above for front location (**S5.3**) and population size (**S5.4**). We simply adjust the replicate-level random effect variance-covariance matrix to (1) link the two model (2) make it treatment-specific, in order to obtain treatment-specific correlations (\(i\) denotes the replicate, \(j\) the treatment, and parameter names are as in **S5.3** and **S5.4**):

\[
\begin{bmatrix}
\alpha\_{i,j} \\
\gamma\_{i,j} \\
\zeta\_{i,j} \\
\eta\_{i,j} \\
\end{bmatrix} \sim
\mathrm{MVNormal}(
\begin{bmatrix}
0\\
0\\
0\\
0\\
\end{bmatrix},\boldsymbol{\Omega}\_{j}),
\]

with \(\boldsymbol{\Omega}\_{j}\) the now treatment-specific covariance matrix and \(\boldsymbol{\mathrm{R}}\_{j}\) the corresponding correlation matrix:

\[
\boldsymbol{\Omega}\_{j} =
\begin{bmatrix}
\sigma\_{\alpha[j]} & 0 & 0 & 0 \\
0 & \sigma\_{\gamma[j]} & 0 & 0 \\
0 & 0 & \sigma\_{\zeta[j]} & 0 \\
0 & 0 & 0 & \sigma\_{\eta[j]} \\
\end{bmatrix}
\times \boldsymbol{\mathrm{R}}\_{j} \times
\begin{bmatrix}
\sigma\_{\alpha[j]} & 0 & 0 & 0 \\
0 & \sigma\_{\gamma[j]} & 0 & 0 \\
0 & 0 & \sigma\_{\zeta[j]} & 0 \\
0 & 0 & 0 & \sigma\_{\eta[j]} \\
\end{bmatrix}.
\] All other parameters and all priors are as in the corresponding univariate models.

#### S5.6 - validation of computer estimates of population sizes (Supplementary Material S2)

To compare computer estimates of parasitism rate \(P\_{i,m}\) with human counts of parasitised eggs \(n\_{i}\) (where \(i\) is a given patch and \(m\) is the estimating computer macro), we used the following bivariate mixed model:

\[
n\_{i} \sim \mathrm{Binomial}(N = 450, p\_{i}), \\
P\_{i,m} \sim \mathrm{Beta}(\mu\_{i}, \phi), \\
\mathrm{logit}(p\_{i}) = \beta\_{0} + \alpha\_{i} ,\\
\mathrm{logit}(\mu\_{i}) = \beta\_{1} + \gamma\_{i},
\] in which the patch-level random effects \(\alpha\) and \(\gamma\) are correlated and distributed as follows:

\[
\begin{bmatrix}
\alpha\_{i} \\
\gamma\_{i} \\
\end{bmatrix} \sim
\mathrm{MVNormal}(
\begin{bmatrix}
0\\
0\\
\end{bmatrix},\boldsymbol{\Omega}), \\
\boldsymbol{\Omega} =
\begin{bmatrix}
\sigma\_{\alpha} & 0 \\
0 & \sigma\_{\gamma} \\
\end{bmatrix}
\times \boldsymbol{\mathrm{R}} \times
\begin{bmatrix}
\sigma\_{\alpha} & 0 \\
0 & \sigma\_{\gamma} \\
\end{bmatrix},
\] where \(\Omega\) and \(\boldsymbol{\mathrm{R}}\) again refer the the covariance and correlation matrices, respectively. As in similar models described above, we use a \(\mathrm{Normal}(0,1.5)\) for fixed effect (\(\beta\)) parameters, a \(\mathrm{Half\mbox{-}Normal}(0,1)\) prior for standard deviations \(\sigma\) and for \(1/\phi\), and a \(\mathrm{LKJCorr}(2)\) prior for the correlation matrix \(\boldsymbol{\mathrm{R}}\).

We ran a second version of this model, to determine whether computer macros presented systematic estimation biases. In this case, the parameter \(\beta\_{1}\), which denotes the (logit-transformed) average parasitism rate in the “computer estimates” model, was simply changed to be macro-specific: \[
P\_{i,m} \sim \mathrm{Beta}(\mu\_{i,m}, \phi), \\
\mathrm{logit}(\mu\_{i,m}) = \beta\_{1[m]} + \gamma\_{i},
\] with the other parts of the model remaining unchanged.

#### S5.7 - Validation of the experimental design (Supplementary Material S3)

As a measure of dispersal distance, we used the mean distance from release \(\bar{D}\_{i,m}\) between the eggs laid in a replicate \(i\) (as observed by macro \(m\)) and the corresponding parental patch using data from the starting generation, when there were only 5 patches per replicate (including the release patch). This means \(\bar{D}\_{i,m}\) can only take values in the \(\left]0, 4\right[\) range, and can be modelled as follows:

\[
\frac{\bar{D}\_{i,m}}{4} \sim \mathrm{Beta}(\mu\_{i,m}, \phi), \\
\mathrm{logit}(\mu\_{i,m})= \beta\_{\text{TREATMENT}[i]} + \alpha\_{m},\\
\alpha\_{m} \sim \mathrm{Normal}(0,\sigma\_{\alpha}).
\] As in other Beta models, we used a \(\mathrm{Normal}(0,1.5)\) prior for the treatment-specific intercepts \(\beta\), and \(\mathrm{Half\mbox{-}Normal}(0,1)\) priors for both the random effect standard deviation \(\sigma\_{\alpha}\) and \(1/\phi\).

### S6 - Supplementary figures for main text models (simulated data)

For simulations, it can be useful to show a bit more precisely how well the theory-based model(s) fit the simulated data. (For experimental data, there are already plots filling that role in the main text)

#### Expansion velocity

Looking at front location as a fonction of time will not make for informative figures, since the overall linear pattern will mask any fine differences. Let’s look instead at the front velocity and how it converges:

**Figure S6.1** Observed and predicted expansion velocities (distance to the starting patch divided by time since start) as a function of \(K\), time since start and treatment. Observed simulation data are summarised by their means and 2.5/97.5% quantiles to keep the figure readable. The mean prediction curves are displayed along with their 95% credible bands. Note how predictions based on the power-law model start to systematically underestimate true speeds towards the end of the simulations, while the exponential predictions catch the fact speed has converged.

#### Genetic diversity

**Figure S6.2** Observed (colored lines) and predicted (grey bands) among-replicate genetic variances in simulations as a function of \(K\), time since start and treatment. The mean prediction curves are displayed along with their 95% credible bands.

### S7 - Summary tables for main text models

We here present posterior means and 95% credible intervals for some parameters of interest of the models discussed in the main text. For conciseness, we:

- focus on fixed effect parameters as they are the ones that interest us in the present study;
- only present the information from the model actually discussed in the main text when there are two competing models (i.e. exponential vs. power for velocities).

The interested reader can get the same information for all other parameters by running the corresponding `Rmd` scripts (scripts 1 and 3 in archived code), which contain various summary calls. Parameter names in the tables below are as in **Supplementary Material S5**.

#### Expansion velocity

**Table S7.1** Posterior means [95% credible intervals] for the model fitted to analyse expansion speeds of *simulated* range expansions

|  | reference | density-dependent dispersal | weak Allee effect | reduced connectivity | reduced connectivity + DDD |
| --- | --- | --- | --- | --- | --- |
| \(\log\_{10}(\lambda)\), \(K\)=225 | 0.08 [0.03, 0.12] | -0.11 [-0.16, -0.07] | 0.14 [0.09, 0.18] | 0.34 [0.3, 0.39] | 0.24 [0.19, 0.29] |
| \(\log\_{10}(\lambda)\), \(K\)=450 | -0.02 [-0.07, 0.03] | -0.21 [-0.26, -0.16] | 0.02 [-0.03, 0.06] | 0.26 [0.22, 0.31] | 0.13 [0.09, 0.18] |
| \(\mathrm{logit}(v\_{t \to \infty})\), \(K\)=225 | 1.35 [1.34, 1.37] | 1.81 [1.8, 1.83] | 0.91 [0.89, 0.92] | 0.56 [0.55, 0.58] | 0.94 [0.92, 0.95] |
| \(\mathrm{logit}(v\_{t \to \infty})\), \(K\)=450 | 1.46 [1.44, 1.47] | 1.87 [1.85, 1.89] | 1.09 [1.08, 1.1] | 0.66 [0.65, 0.67] | 0.98 [0.96, 0.99] |

**Table S7.2** Posterior means [95% credible intervals] for the model fitted to analyse expansion speeds of *experimental* range expansions

|  | reference | reduced connectivity |
| --- | --- | --- |
| \(\log\_{10}(\lambda)\) | -0.39 [-0.64, -0.16] | -0.61 [-0.93, -0.31] |
| \(\log(v\_{t \to \infty})\) | -0.03 [-0.19, 0.14] | -0.07 [-0.25, 0.11] |
| \(\log(v\_{t = 1})\) | 0.71 [0.43, 0.96] | 0.63 [0.37, 0.87] |

#### Genetic diversity

**Table S7.3** Posterior means [95% credible intervals] for the model fitted to analyse genetic diversity of *simulated* range expansions

|  | reference | density-dependent dispersal | weak Allee effect | reduced connectivity | reduced connectivity + DDD |
| --- | --- | --- | --- | --- | --- |
| \(\log\_{10}(\lambda)\), \(K\)=225, core patches | -2.69 [-2.71, -2.67] | -2.82 [-2.84, -2.79] | -2.67 [-2.69, -2.65] | -2.53 [-2.54, -2.51] | -2.69 [-2.71, -2.67] |
| \(\log\_{10}(\lambda)\), \(K\)=450, core patches | -2.97 [-3, -2.94] | -2.97 [-3, -2.95] | -2.93 [-2.96, -2.91] | -2.95 [-2.98, -2.92] | -3.07 [-3.1, -3.04] |
| \(\log\_{10}(\lambda)\), \(K\)=225, edge patches | -0.89 [-0.91, -0.86] | -1.22 [-1.25, -1.2] | -1.03 [-1.05, -1] | -0.78 [-0.81, -0.74] | -1.06 [-1.09, -1.04] |
| \(\log\_{10}(\lambda)\), \(K\)=450, edge patches | -0.96 [-0.99, -0.93] | -1.41 [-1.42, -1.39] | -1.25 [-1.27, -1.23] | -0.89 [-0.92, -0.86] | -1.23 [-1.25, -1.2] |

|  | all treatments |
| --- | --- |
| \(\mathrm{logit}(V\_{max})\) | -1.12 [-1.13, -1.12] |

**Table S7.4** Posterior means [95% credible intervals] for the model fitted to analyse genetic diversity of *experimental* range expansions

|  | reduced connectivity | reference |
| --- | --- | --- |
| average \(\log\_{10}\)(decay rate of genetic diversity) (\(\beta\_{1}\)) | -1.65 [-1.79, -1.5] | -1.66 [-1.84, -1.49] |
| difference edge-core in decay rate (\(\beta\_{2}\)) | 0.3 [0.14, 0.46] | 0.72 [0.5, 0.98] |

|  | all treatments |
| --- | --- |
| logit of initial genetic diversity (\(\beta\_{0}\)) | -0.63 [-0.7, -0.56] |

#### Population size (experimental data)

**Table 7.5** Posterior means [95% credible intervals] for the *univariate* model fitted to analyse mean population size in core patches (x = 0) during *experimental* range expansions

|  | reference | reduced connectivity |
| --- | --- | --- |
| logit of carrying capacity (\(K\)) | 0.4 [-0.22, 0.98] | 0.57 [-0.03, 1.16] |
